# Tetrameric UvrD helicase is located at the *E. coli* replisome due to frequent replication blocks

**DOI:** 10.1101/2021.02.22.432310

**Authors:** Adam J. M Wollman, Aisha H. Syeda, Jamieson A. L. Howard, Alex Payne-Dwyer, Andrew Leech, Dominika Warecka, Colin Guy, Peter McGlynn, Michelle Hawkins, Mark C. Leake

## Abstract

DNA replication in all organisms must overcome nucleoprotein blocks to complete genome duplication. Accessory replicative helicases in *Escherichia coli*, Rep and UvrD, help remove these blocks and aid the re-initiation of replication. Mechanistic details of Rep function have emerged from recent live cell studies; however, the division of UvrD functions between its activities in DNA repair and role as an accessory helicase remain unclear in live cells. By integrating super-resolved single-molecule fluorescence microscopy with biochemical analysis, we find that UvrD self-associates into tetrameric assemblies and, unlike Rep, is not recruited to a specific replisome protein despite being found at approximately 80% of replication forks. Instead, its colocation with forks is likely due to the very high frequency of replication blocks composed of DNA-bound proteins, including RNA polymerase and factors involved in repairing DNA damage. Deleting *rep* and DNA repair factor genes *mutS* and *uvrA*, and inhibiting transcription through RNA polymerase mutation and antibiotic inhibition, indicates that the level of UvrD at the fork is dependent on UvrD’s function. Our findings show that UvrD is recruited to sites of nucleoprotein blocks via different mechanisms to Rep and plays a multi-faceted role in ensuring successful DNA replication.

## INTRODUCTION

Replication and transcription of DNA occur simultaneously in *Escherichia coli,* making conflicts between the bacterial replisome, the molecular replication machine comprising in excess of 10 different proteins, and RNA polymerase (RNAP) inevitable (1–4). Collisions between replication and transcription machineries hinder replication fork progression and cause genome instability (5–14). RNAP can pause, stall and backtrack while actively transcribing, and any immobile RNAP presents long-lived barriers to replisomes (15, 16); RNAP is the most common nucleoprotein obstacle for translocating replisomes. DNA is also frequently damaged in normal growth conditions due to exposure to endogenous and exogenous DNA damaging agents, which also presents significant obstacles for replication (17–19).

Several mechanisms exist to reduce conflict between the replisome and nucleoprotein blocks. For example, transcription elongation and termination factors reduce the number of immobile RNAPs on DNA (1, 18, 20–24). Head-on collisions were originally thought to be more harmful (25–27) and therefore more important to resolve; a view supported by the occurrence of highly transcribed genes encoded on the leading strand, presumably to ensure co-directional transcription and replication (26, 28, 29). However, co-directional collisions of the replisome and RNAP also impact fork progression (3, 18, 30).

Accessory replicative helicases, identified in prokaryotes and eukaryotes, promote fork movement through resolving nucleoprotein blocks (1, 12, 13, 31, 32). The Rep and UvrD accessory helicases in *E. coli*, as well as the Pif1 helicase in *Saccharomyces cerevisiae*, perform important roles in helping to clear nucleoprotein complexes (33). More recently, Pif1 has been proposed to be a general displacement helicase for replication bypass of protein blocks, a function similar to the Rep and UvrD helicases (34). These helicases can reduce replisome pausing at several types of native transcription complexes in living cells (13, 22, 35), and of engineered nucleoprotein blocks in biochemical experiments *in vitro* (1, 31).

Rep is the primary accessory helicase in *E. coli*, however, UvrD, a superfamily 1 helicase that translocates 3’-5’ along ssDNA, partially compensates for its absence *in vivo,* attributed to the high degree of homology between these two helicases and the abundance of UvrD inside cells (12, 21, 36–38). Rep also promotes replication through stalled transcription elongation complexes (1, 12, 13, 39, 40). UvrD’s role in nucleotide excision repair along with UvrABC proteins, as well as in methyl-directed mismatch repair along with the MutHLS proteins, is well characterised (41–46). UvrD interacts directly with RNAP but the functional importance of this interaction has been unclear until recently (47–51). UvrD has been suggested to promote backtracking of stalled RNAP as a first step in transcription-coupled repair (50, 52); however, other studies argue against UvrD playing any role in coupling nucleotide excision repair to stalled transcription complexes (48, 49, 53–57).

Our aim in this study was to establish the role of UvrD during DNA replication. The primary objective was to test whether UvrD is present at active replication forks using dual-color single-molecule imaging of fluorescently labelled UvrD and replication forks in living *E. coli* cells. We conclude that UvrD is present at replication forks but is not recruited via specific protein interactions with the replisome, which means that it operates in a fundamentally different way to that of Rep. By using super-resolved millisecond single-molecule fluorescence microscopy of mGFP-UvrD in live cells co-expressing a fork marker, DnaQ-mCherry, we show that 80% of DnaQ foci colocalise with UvrD. Stepwise photobleaching intensity analysis reveals tetrameric UvrD foci *in vivo*, to be compared with earlier observations performed *in vitro* using analytical ultracentrifugation that found a mixture of UvrD dimers and tetramers (58). Surface plasmon resonance (SPR) of UvrD in combination with each replisome component identified no specific replisome protein interaction that would explain replisome-UvrD colocalisation. Deletion of Rep, mismatch repair protein MutS and nucleotide excision repair protein UvrA, and disruption of transcription using either mutated RNAP or antibiotic inhibition, significantly reduces the number of UvrD or level of observed colocalisation of UvrD with the fork. Coupled with *in vitro* block-dependent differences in UvrD replication promotion efficiency, we conclude that the colocation of UvrD with the replisome stems primarily from UvrD resolving frequent nucleoprotein blocks to the DNA replication machinery, as opposed to it binding specifically to a replisome component, contrasting distinctly with the replisome-binding mode of operation of Rep.

## RESULTS

### UvrD is present at the majority of replication forks

We investigated UvrD’s role in DNA replication using super-resolved single-molecule fluorescence microscopy to track dynamic patterns of fluorescently labelled UvrD localization relative to the replication fork in live cells (Figure 1A). We performed dual-color Slimfield (59, 60) to assess the degree of colocalisation between UvrD and the replisome, using the same imaging protocol used previously to confirm Rep colocalisation with the replisome (13). We employed DnaQ-mCherry as a fork marker-DnaQ encodes a core epsilon subunit of DNAP (13, 61–64)-along with genomically integrated mGFP-UvrD to report on the localization of UvrD. These strains phenocopied wild type for growth and in plasmid loss assays for mGFP-UvrD function retention (Supplementary Figures S1, S2, Tables S1-4). *In vitro* unwinding assays indicate that the nucleoprotein block removal activity of mGFP-UvrD was comparable to wild type (Supplementary Figure S3; note, a summary of all activity comparisons between mGFP-UvrD and wild type UvrD is shown in Table S5). Using Slimfield, we observed 1-2 DnaQ foci per cell, as well as a background ‘pool’ of unbound DnaQ molecules in the cytoplasm (61), as in the previous study for Rep, corresponding to the two moving forks which appear as a single focus near the start of replication when they are separated by less than the 200-300 nm diffraction limit of our microscope (Figure 1A,B Supplementary Figure S4). We also observed 1-2 UvrD foci per cell (and associated pool), of which 67 ± 7% (±SEM) were colocalised with DnaQ foci and 84 ± 5% of DnaQ foci colocalised with UvrD (Figure 1B and C). We calculated the probability of random overlap between foci by modelling nearest-neighbor separation as a Poisson distribution (65), indicating 20%. To confirm that colocalisation between DnaQ and UvrD was not just a function of the nucleoid association of UvrD, we performed Slimfield on Heat-stable nucleoid-structuring protein (H-NS) tagged with GFP-H-NS associates with the nucleoid but not specifically with the fork-and co-labelled DnaQ-mCherry. We found similar numbers of H-NS foci to UvrD (Supplementary Figure S5) but only 38 ± 4% DnaQ foci were colocalised with H-NS foci implying that, similar to Rep (13), UvrD is present at the majority of forks.

**Figure 1:**
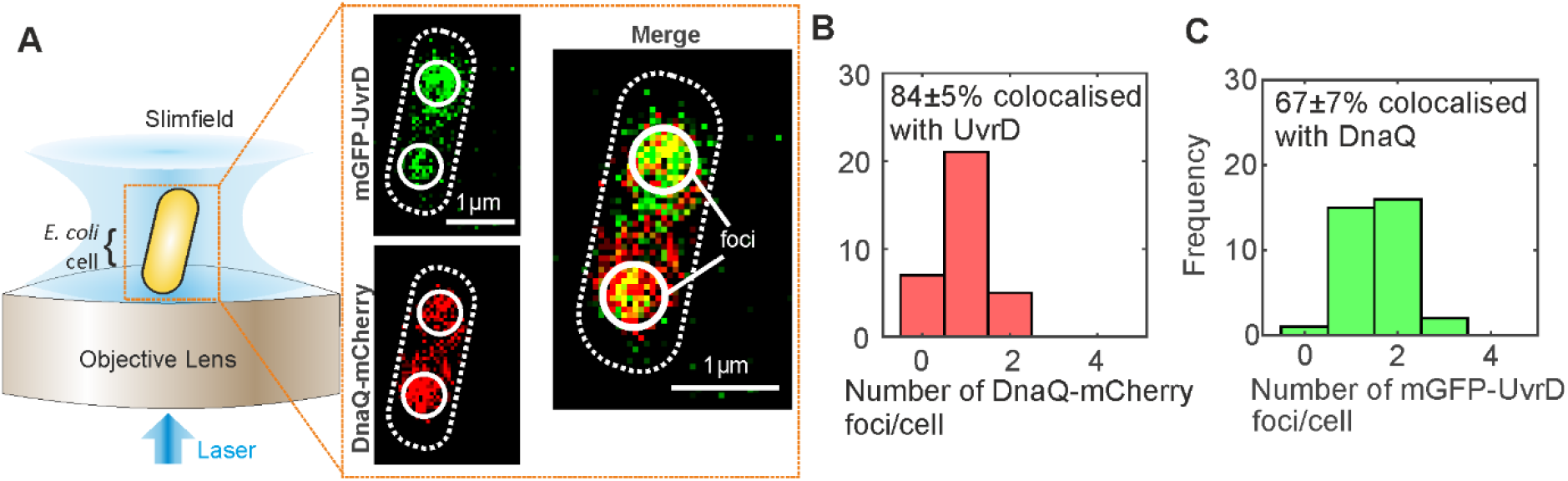
Super-resolved single-molecule light microscopy indicates fork-UvrD colocation. (A). Slimfield microscope schematic and micrographs of mGFP-UvrD, DnaQ-mCherry. Cell membrane and DnaQ/UvrD foci indicated as white dashed and non-dashed lines respectively. (B) and (C) Histogram showing the number of DnaQ and UvrD foci detected per cell respectively, SD errors. N=45 cells.

### UvrD foci are tetrameric assemblies in live bacteria

Slimfield also enabled the stoichiometry of these foci (i.e., the number of fluorescently-labelled molecules within each focus) to be determined by utilizing a method involving stepwise photobleaching analysis (61). Here, we determined the brightness of a single-molecule of mGFP or mCherry fluorescent protein using Slimfield (Supplementary Figure S7), and used these values to normalize the initial brightness of tracked foci to estimate the stoichiometry. We found that DnaQ had the same range of 2-6 molecules per focus (Supplementary Figure S8) as found previously, corresponding to 2-3 polymerases per fork (13, 61). The distribution of the stoichiometry of UvrD foci which are colocalised with DnaQ had a distinct lowest-order peak corresponding to approximately four molecules, with subsequent peaks at multiples of four (Figure 2A). UvrD foci that were not colocalised with DnaQ also contained a distinct lowest-order peak at four, but subsequent peaks were less clearly tetrameric.

**Figure 2.**
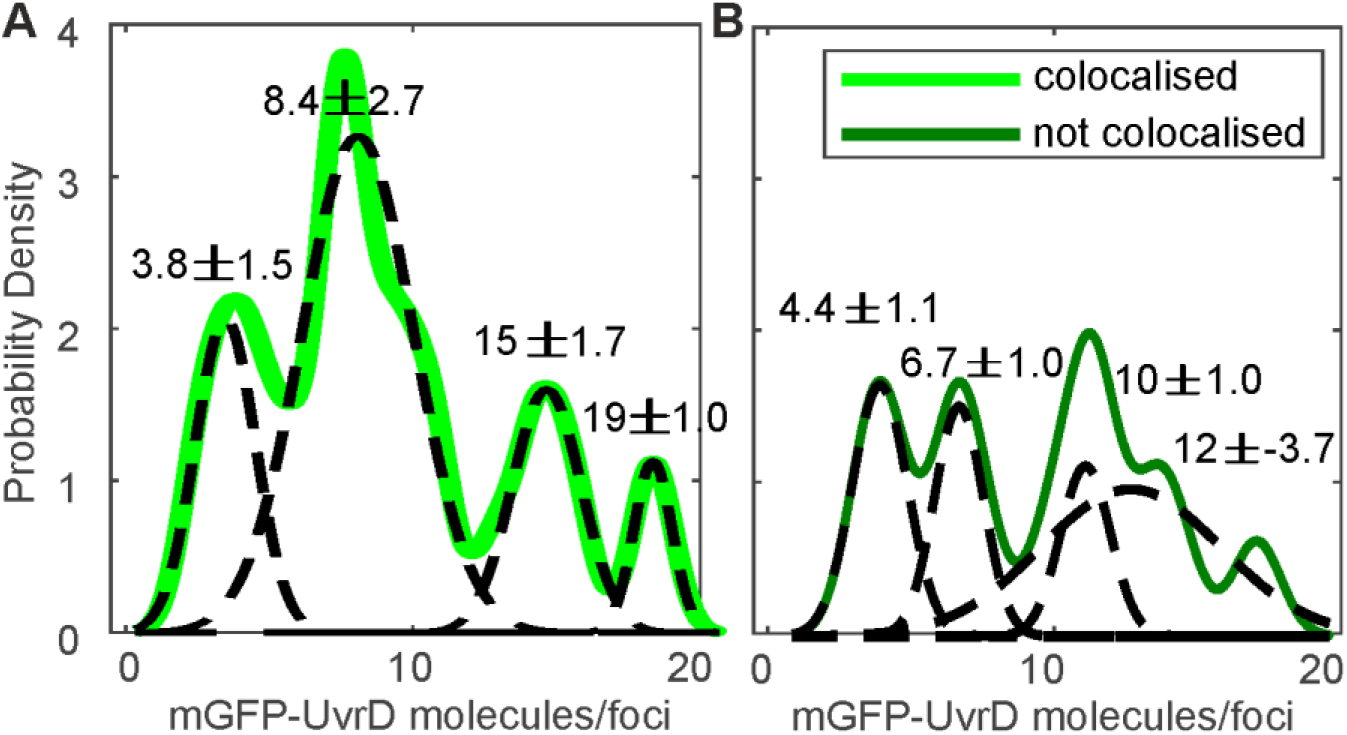
UvrD stoichiometry shows tetrameric periodicity. The distribution of UvrD foci stoichiometry rendered as a kernel density estimate for foci colocalised (A) and not colocalized (B) with DnaQ foci. Distributions were fitted with multiple Gaussian peaks (black dotted lines) with the peak value ± error determined by the half width at half maximum indicated above each peak

Although we utilised a definitive monomeric variant of GFP which contained an A206K mutation to inhibit dimerization (66), we did not want to dismiss possible non-physiological influences of fluorescent protein labelling of UvrD in its self-assembly process. Using Size Exclusion Chromatography Multi-Angle Laser Light Scattering (SEC-MALLS) we found qualitatively that the majority of UvrD runs off in a monomeric fraction for both purified wild type UvrD and for mGFP-UvrD (Supplementary Figure S9). However, SEC-MALLS has known limitations in differentiating between monomeric and oligomeric fractions due to non-specific binding to the SEC resin (depending on buffering conditions), resulting in proteins eluting later than expected and co-elution of monomeric species and higher mass oligomers. In addition, there are sensitivity difficulties for detecting components present in relatively small quantities – we estimate the error on the cumulative weight fraction to be ∼20% on our system, consistent with the reported sensitivity error reported by others (67, 68).

To overcome these limitations, we performed high-precision high-sensitivity Slimfield of purified mGFP-UvrD conjugated to a glass coverslip via anti-GFP antibody (69) enabling detection of fluorescent foci bound to the surface with single-molecule sensitivity. Slimfield indicated between ∼10-20 surface-bound foci per field of view (Supplementary Figure S9A), whose 200-300 nm width (half width at half maximum, HWHM) was comparable to that determined for single purified GFP molecules. We measured the fluorescence brightness of each focus using the same automated single-particle tracking software as for Slimfield *in vivo* (Supplementary Figure S9B), whose tracks photobleached in a stepwise manner consistent with the presence of just a single GFP molecule. The amplitude of the probability distribution function of foci brightness values (Supplementary Figure S9B) indicated that the proportion of dimers (corresponding to a brightness of approximately twice the modal value) is ∼3% that of the corresponding value for monomers. Although it is not possible to entirely exclude non-physiological effects of any fluorescent protein on the cellular activities of a tagged protein, the finding from this single-molecule assay shows that the proportion of monomeric mGFP-UvrD is substantially higher than dimeric mGFP-UvrD and so, if there is an influence of mGFP on the self-assembly process of UvrD, it is unlikely to be a dominant factor.

At higher concentrations, UvrD has been observed to exist in a monomer-dimer-tetramer equilibrium *in vitro* (58). DNA unwinding experiments *in vitro* suggest UvrD must function as at least a dimer (70) or a trimer (71), however the oligomeric state of functional UvrD is disputed (reviewed in (72)). Our results are consistent with UvrD existing predominantly as a tetramer, or a dimer of dimers, with multiple tetramers or dimer-dimers sometimes present at forks. It should be noted that we often observe only a single DnaQ focus corresponding to two forks which are separated by less than the diffraction limit of our microscope (Supplementary Figure S8) such that some multiple tetrameric UvrD foci correspond to two replisomes. The less distinct peaks in the UvrD foci that are not colocalised with DnaQ may indicate that other UvrD species (monomers, dimers or trimers) are also present.

We determined the total cell copy number of UvrD by estimating the total contribution of mGFP-UvrD fluorescence to the observed intensity in whole cells (73), indicating approximately 800 molecules of UvrD per cell (Supplementary Figure S10), similar to that reported previously for Rep (13), and in line with other proteins that have a small active population detected as foci and a larger population of diffusive molecules in a protein pool (61, 73–76). This is a lower level of UvrD than was reported using transcription rate measurements (77) or high-throughput proteomics (78) but is comparable to recent estimates using ribosome profiling (79).

### There is no obvious replisome interaction partner for UvrD at the fork

Our earlier findings concerning Rep indicated that colocalisation with the fork and recruitment to the replisome were mediated by protein-specific interaction with the replicative helicase DnaB (80), consistent with Rep and DnaB both exhibiting hexameric stoichiometry (13, 61). We sought to determine whether UvrD is also recruited by interacting with a specific replisome component, expanding on a previous study which used SPR to test whether purified UvrD interacted with each replisome component(12). In this study, UvrD, under conditions consistent with predominantly monomeric UvrD (Supplementary Figure S9), exhibited no direct interaction with primase, SSB, β sliding clamp, the DNA polymerase III αɛ, χψ or γ complexes; or θ, δ, δ′, χ, and γ subunits (12). However, it did identify a putative weak UvrD-Tau interaction which we interrogated further with more sensitive SPR experiments. We were able to replicate the weak UvrD-Tau interaction; however, as a control we tested Rep and found this also interacted with Tau to a similar extent (Supplementary Figure S11). Tau interactions with either helicase were ∼100x lower affinity than UvrD with known interaction partner UvrB (Supplementary Figure S11). We therefore conclude that the Tau interaction with UvrD is likely to be non-specific and that, unlike Rep, UvrD does not have an obvious, direct interaction partner at the replisome, at least from the suite of replisome proteins tested. Since our findings suggest that UvrD may exist as as a tetramer at the replisome, it is also unlikely to be recruited by Rep, which has shown putative interaction (81).

### The presence of UvrD at the fork is mediated by specific replication blocks

Alternatively, we hypothesized that the association of UvrD with the replication fork might be dependent on its activity in resolving nucleoprotein blocks. To test, we systematically impaired key DNA replication repair processes in which UvrD is implicated, as well as perturbing RNAP replicative blocks by disruption of transcription through both genetic mutation and antibiotic inhibition (Figure 3). We investigated how these perturbations altered fork association through UvrD colocalisation with DnaQ. As a control, we compared these to Rep colocalisation with DnaQ which can be abrogated down to random levels of colocalisation, indicating no interaction with the fork, by mutations which stop Rep interacting with DnaB and PriC (Figure 3A). We perturbed UvrD’s activity in resolving DNA damage at replication blocks by deleting *mutS* or *uvrA*, rendering the strains defective in UvrD-associated mismatch repair or nucleotide excision repair respectively (43, 82). We reduced the need for UvrD to remove transcriptional blocks to replication (49) by introducing the *rpoB*35* mutation, which destabilizes open complexes formed during initiation of transcription (83, 84); and by using the antibiotic rifampicin, which blocks elongation of RNA transcripts (85). All of these strains were healthy under normal growth conditions (Supplementary Figure S1, Table S4) and displayed the same phenotype for DnaQ foci number and stoichiometry within measurement error (Supplementary Figure S8).

**Figure 3:**
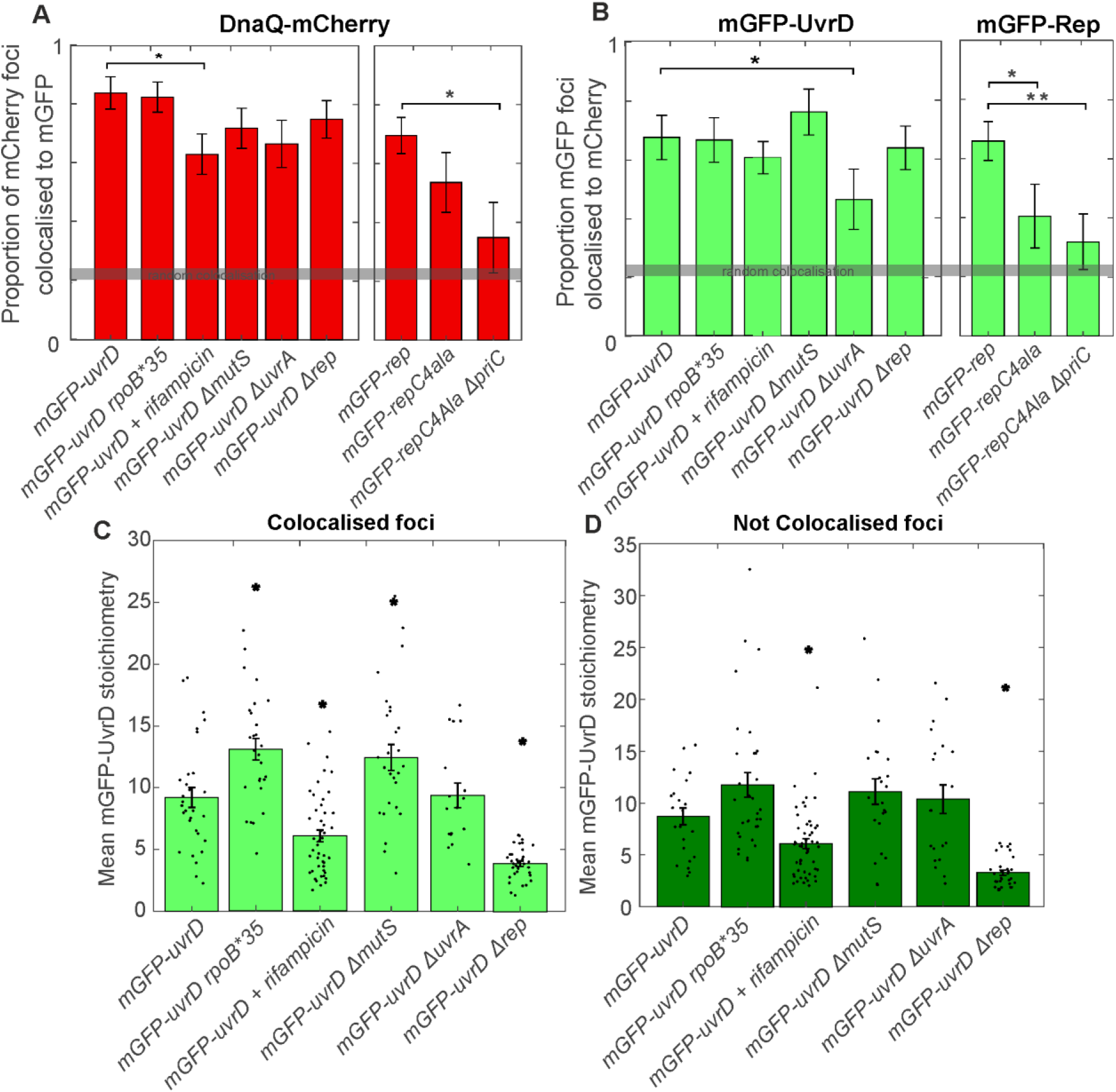
**Perturbing UvrD function using mutations and rifampicin antibiotic intervention influences fork-UvrD colocation**. A. and B. The mean proportion of colocalised DnaQ and UvrD or Rep foci per cell respectively. Error bars represent standard error in the mean and statistical significance by Brunner-Munzel and Student’s t-test p<0.05 indicated by *. Gray bar indicates levels of predicted random colocalisation. C. and D. Jitter plots of colocalised and not colocalised UvrD stoichiometry respectively. Bars show mean UvrD stoichiometry with error bars showing the standard error in the mean and statistical significance by Brunner-Munzel and Student’s t-tests p<0.05 indicated by *. N=30 cells.

For the transcriptional block perturbations, the *rpoB*35* mutant dual labelled strain exhibited no difference in the proportion of colocalised DnaQ or UvrD foci (Figure 3A and B). This mutant and all others tested exhibited the same predominant tetrameric stoichiometry trends as wild type (Supplementary Figure S12), with clear peaks at tetramer intervals in the colocalised stoichiometry distribution and less clear but still largely tetrameric intervals in the stoichiometry distribution for UvrD foci that are not colocalised with DnaQ. This provides further evidence for tetrameric UvrD assemblies *in vivo*. We compared the mean stoichiometry for each mutant to that of wild type (Figure 3C and D). Unexpectedly, the *rpoB*35* mutant produced a statistically significant increase in mean colocalised stoichiometry, corresponding to approximately one extra UvrD tetramer at the replication fork (Figure 3C). This finding may indicate the level of instability conferred by this mutation has multiple downstream effects that alter UvrD localisation significantly at the fork. Alternatively, it could mean that *rpoB*35* does not affect the formation of the types of transcriptional block that are acted upon by UvrD. Rifampicin treatment produced a clearer response, reducing the number of DnaQ foci colocalised with UvrD by ∼20% (Figure 3A) and the mean stoichiometry of colocalised and not colocalised UvrD foci by around one UvrD tetramer (Figure 3C and D). Single-molecule observations of RNAP have shown it to be significantly more mobile under rifampicin treatment (86), implying fewer RNAPs bound to DNA and fewer blocks to replication. Our rifampicin treatment results are consistent with fewer transcriptional blocks to replication reducing the amount of UvrD recruited to the replisome and support the hypothesis that UvrD is recruited by these replicative blocks.

By deleting *mutS* or *uvrA* we removed UvrD’s role in mismatch repair and nucleotide excision repair respectively. In the Δ*mutS* mutant we observed no change in UvrD colocalisation with DnaQ, but did measure an increase in the mean UvrD stoichiometry colocalised with DnaQ by around one tetramer (Figure 3A). In the Δ*uvrA* mutant we observed a decrease of ∼25% in UvrD colocalisation with DnaQ (Figure 3B) with no change in the number of UvrD foci (Supplementary Figure S8). This agrees with a recent model that UvrA and UvrD are continuously associated with a subpopulation of RNAP to form a pre-transcription-coupled-repair complex (pre-TCRC); UvrD is then recruited at DNA lesions to facilitate DNA repair (50).

UvrD has been shown to compensate for Rep in *rep* null mutants (12). We also tested what effect this mutation would have in Slimfield *in vivo*. Surprisingly, *rep* deletion resulted in a steep drop in UvrD copy number (Supplementary Figure S10). The degree of colocalisation between UvrD and DnaQ remained unchanged compared to wild type (Figure 3A and B) but foci stoichiometry was reduced to a single UvrD tetramer (Figure 3C and D and Supplementary Figure 11), consistent with the observed lower copy number and availability of UvrD. The drop in UvrD copy number may reflect complex regulatory shifts in response to losing a key protein in Rep, although it is interesting that even at low copy numbers, a UvrD tetramer remains recruited to the fork. Double knockouts, deficient in both Rep and UvrD, are lethal (12) so this minimal helicase presence must be the limit required for survival.

### Dynamic association of UvrD with the fork depends on DNA repair processing

By tracking UvrD foci, we were also able to characterize their dynamics in wild type and repair impaired cells. We measured the exposure time per frame multiplied by the number of image frames in which UvrD foci were colocalised with DnaQ foci and fitted the distribution of these dwell times with a single exponential function. We also calculated the apparent microscopic diffusion coefficient of individual foci (Figure 4 and Supplementary Figure S13). The mean UvrD dwell time at the fork (corresponding to the exponential fit time constant) for the wild type strain was measured to be approximately 4 ms, indicating a relatively dynamic association of UvrD at the replication fork. The diffusion coefficients (*D*) were best fit with a three parameter Gamma model, containing immobile (*D* = 0.1 µm^2^/s), transiently immobile population (*D* = 0.5 µm^2^/s) and mobile (*D* = 1.2 µm^2^/s) populations similar to Rep. In wild type, we found that approximately 10% of UvrD foci were immobile, whether colocalised with the fork or not, with the remaining foci split between the other two populations.

**Figure 4:**
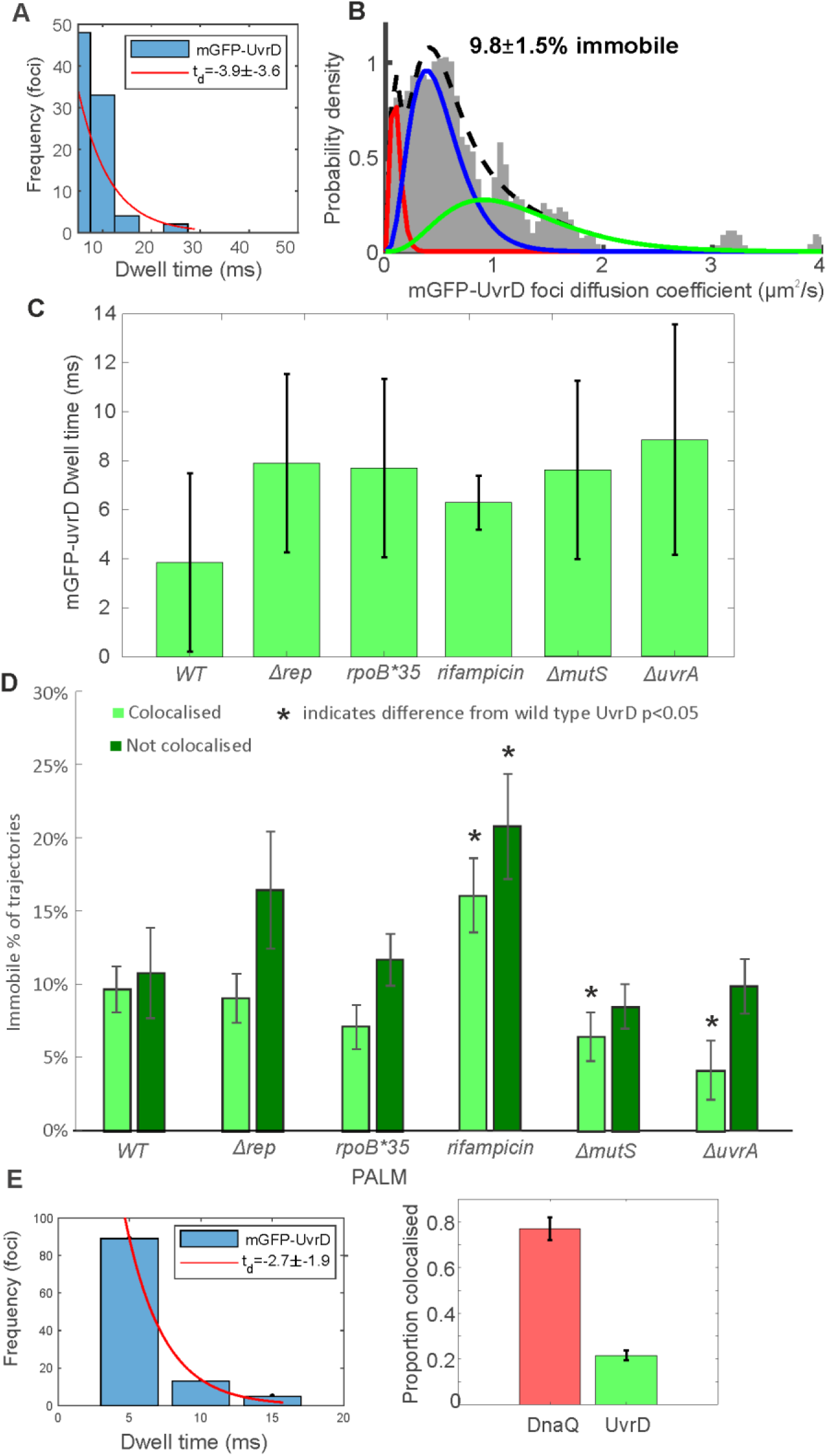
UvrD dynamics are heterogeneous. A. Distribution of time over which UvrD and DnaQ foci were colocalised (blue) fitted with an exponential (red) to yield a characteristic dwell time. B. Distribution of microscopic diffusion coefficients of wild type UvrD foci (gray) fitted with a three-state gamma distribution model containing a relatively immobile population with *D* = 0.1 µm^2^/s (red), a transiently immobile population, *D* = 0.5 µm^2^/s (blue) and a mobile population, D=1.2 µm^2^/s (green). C. The dwell times for each UvrD strain. Error bars represent the 95% confidence intervals on the fit. D. The proportion of UvrD foci in the immobile population for each UvrD strain colocalised or not colocalised with DnaQ foci location. Statistically significant differences p<0.05 from wild type indicated by *. E. PALM dwell time and colocalisation proportion of detected tracks. N=45 cells for wild type and 30 per mutant, with ∼300 trajectories.

We applied the same approach to cells which had been perturbed using either a range of mutations which were known to affect DNA replication repair and block resolution, or antibiotic treatment known to affect expression of RNAP. All perturbations resulted in a marginal though statistically insignificant increase in mean dwell times to between 6-9 ms. The Δ*rep* and *rpoB*35* mutations resulted in no change in mobility from wild type. Rifampicin treatment however, resulted in an increase in the proportion of immobile foci both at and away from the fork. This finding appeared at first counterintuitive since rifampicin treatment results in more mobile RNAP (86), however it is unclear what effect rifampicin has on the population of RNAP molecules that remain bound to DNA. These results may suggest that such blocks provide an increased barrier to replication, decreasing UvrD mobility as it attempts to resolve blocks. In Δ*mutS* and Δ*uvrA* strains, the proportion of immobile foci decreased at the fork, again consistent with the hypothesis that UvrD association with the fork is dependent on DNA damage resolution activity. One alternative explanation for these results is that UvrD is less mobile when resolving DNA damage blocks to replication and more mobile when resolving transcriptional blocks, as reducing or eliminating UvrD’s role in these processes decreased or increased the immobile fraction in DNA repair and transcription impaired cells respectively (Figure 4).

### *In vitro* biochemical experiments support the block-dependent UvrD function model

To investigate the role of different nucleoprotein blocks in UvrD recruitment to the replisome, we reconstituted replisomes on plasmid templates containing *oriC* and block sites. This *in vitro* approach allowed us to specify the type of nucleoprotein block and test whether the ability of UvrD to promote replication is affected by whether UvrD recruitment occurs before or after the replisome encounters the block. In these experiments, UvrD or a Rep control were included either from the start of replication, or were added after the replisome encountered the nucleoprotein block. Blocked fork resolution was probed by examining replication products using denaturing gel electrophoresis.

We first generated stalled transcription elongation complexes using the P_lacUV5 52C_ promotor which allows RNAP to be stalled via nucleotide deprivation. The P_lacUV5 52C_ promotor contains no C residues between +1 and +52 on the non-transcribed strand, enabling transcription to be stalled by the omission of CTP. Stalled RNAP acts as a barrier to the replisome in both head-on and co-directional orientations and prevents the formation of full-length replication products (1). Arresting the replisome at this RNAP block resulted in replication products consisting of four truncated leading strands. The strand sizes matched those expected for replisomes moving clockwise or counter-clockwise from *oriC* and encountering P_lacUV5 52C_ or the promoters within the *ColE1* plasmid origin of replication (Figure 5A, lane 2 and Figure 5C) (1). As previously found, inclusion of Rep or UvrD promotes movement of replisomes through stalled RNAP (1). This is shown by a decrease in all four truncated leading strand products and a concomitant increase in production of full-length leading strands (Figure 5A, lanes 3 and 5 and 5E) (1). Addition of UvrD once the replisome had already encountered the block gave variable results that ranged from an inhibitory to a stronger effect, although differences were not statistically significant at p<0.05 (Figure 5A, compare lane 5 with 6, Figure 5E). Similar results were also obtained for Rep (Figure 5A, compare lane 3 with 4, Figure 5E). Neither Rep nor UvrD are capable of pushing stalled RNAP from the P_lacUV5 52C_ stall site in the absence of replication, so these results are dependent on active replisomes encountering blocks (1).

**Figure 5:**
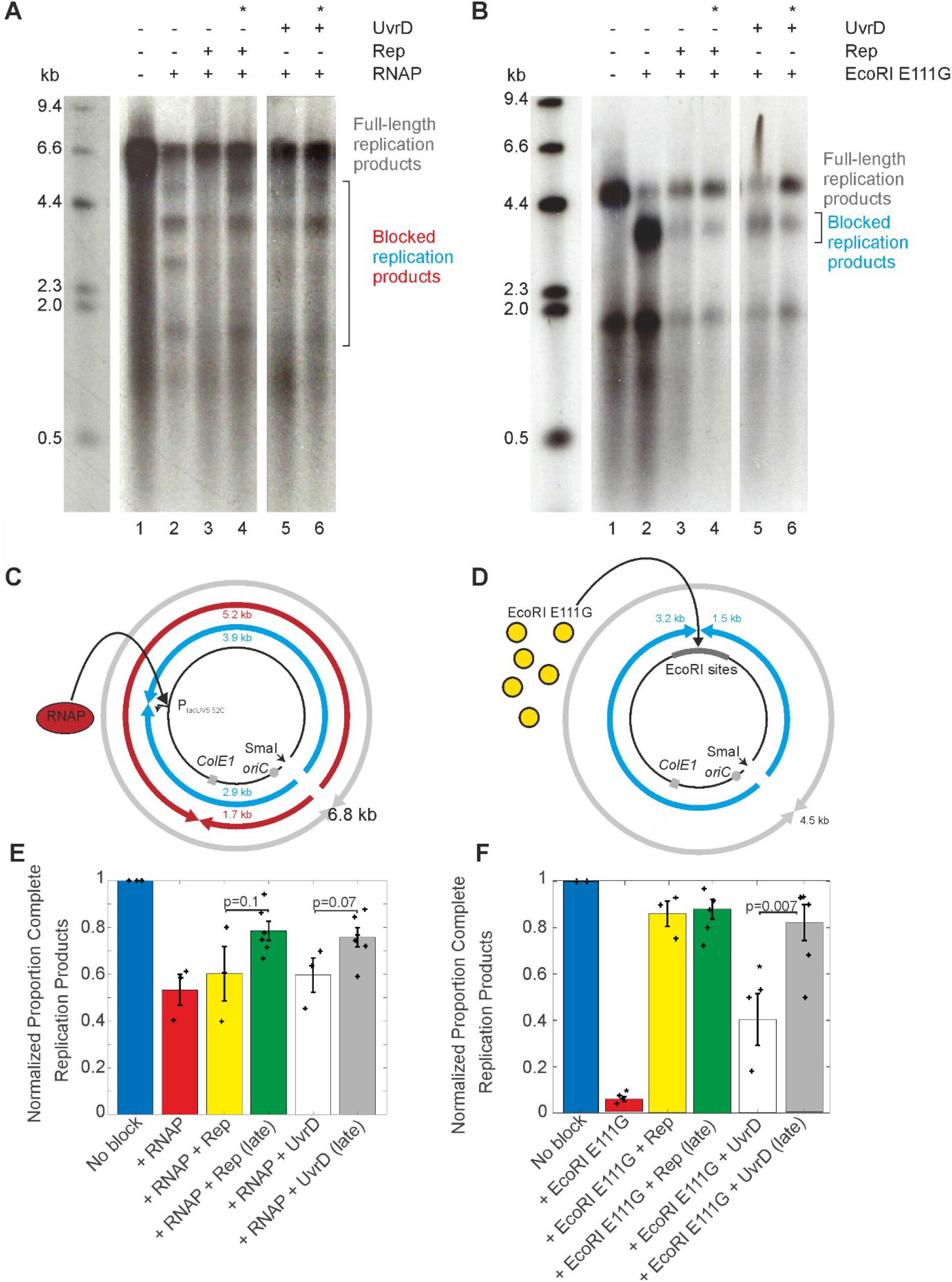
Helicase pre-association affects the promotion of replication fork movement through different nucleoprotein blocks. A. Denaturing agarose gel of replication products formed from pPM872 (1) in the absence and presence of RNAP block, Rep and UvrD added pre-and post-collision (*) as indicated. B. Denaturing agarose gel of replication products formed from pPM594 (12) in the absence and presence of EcoRI E111G block, Rep and UvrD added pre-and post-collision (*) as indicated. Truncated leading strand products formed by replisomes originating from *oriC* and colliding with RNAP (C) and EcoRI E111G (D). Quantification of lanes in (A) and (B) from three technical replicates shown in (E) and (F), with standard error in the mean indicated and each replicate result shown as a cross. Note, we were also able to confirm that the mGFP tag used in the UvrD microscopy experiments does not prevent UvrD from dislodging nucleoprotein blocks in this context (see Supplementary Figure S14).

Since UvrD has been shown to interact with RNAP (47–49), we reasoned that it could be quickly recruited to an RNAP block no matter the time of addition. We used a different nucleoprotein block that does not bind UvrD to test whether UvrD-block pre-association is a requirement for its promotion of replication. Modified restriction enzymes are a useful model for nucleoprotein blocks because their block strength can be modified by varying the number of recognition sites on DNA templates. We chose a mutant restriction enzyme (EcoRI E111G) that binds to its recognition sequence but has reduced cleavage activity (87). EcoRI E111G blocks to replication can be removed by Rep or UvrD helicase activity (12). We tested whether UvrD or a Rep control could promote replication through an array of eight EcoRI sites when added before or after initiation of replication (Figure 5B and D). Rep promotion of replication through this barrier was unaffected by time of addition (Figure 5B, lane 3 and 4, 5F). Presumably this is because the Rep-DnaB interaction ensures it will always be present at replication fork blocks as it travels with the replisome. Surprisingly, adding UvrD post-collision did not impede its promotion of replication; in fact, the resolution of these collisions appeared to be more efficient because the proportion of full-length replication products increased (Figure 5B, compare lane 5 with 6, 5F). Quantification of full-length replication products as a proportion of reaction products showed that late UvrD addition increased full-length replication products to 84% (n=6) compared to 41% (n=3) when UvrD was present before fork collision. This contrasts with the RNAP block results where the UvrD-RNAP interaction (47–50) might be responsible for localising UvrD to the block and facilitating resolution, resulting in no difference when present pre-or post-collision. Similarly, Rep associates directly with the replication fork, resulting in no difference when present pre-or post-collision, for both blocks. These results support the absence of a UvrD-replisome interaction and suggest that UvrD might be recruited to stalled replication forks at nucleoprotein blocks using a different mechanism than Rep. We have no direct evidence to support a hypothesis that UvrD recognises a specific DNA structure at a blocked fork, and note previous studies which indicate no binding specificity to DNA beyond requiring 3ˈssDNA (88, 89). Combining these data with reductions in colocalisation between UvrD and DnaQ fork marker when replication blocks are perturbed *in vivo*, leads us to propose that UvrD is present at the replication fork due to the high frequency of different types of blocks to replication.

## DISCUSSION

Here, we show that most replisomes are associated with UvrD. However, unlike the established Rep association, UvrD has no replisome interaction partner. UvrD instead associates with the replisome through the frequent blocks to replication encountered by the fork. Similarly to Rep, *in vitro* evidence suggests that UvrD functions poorly as a helicase when it is monomeric (90). We showed previously that functional Rep foci are hexameric *in vivo* but that this oligomerisation state is due to interactions with the hexameric DnaB replicative helicase (13). Again, UvrD functions differently and appears to self-associate into tetramers, as evidenced in our findings by tetrameric periodicity both at and away from the fork (Figure 2 and Supplementary Figure 11), consistent with previous ultracentrifugation experiments *in vitro* (58). It is possible that UvrD may not have evolved to fulfil exactly the same functions as Rep; in many enzyme-catalysed reactions, if one enzyme is depleted then another will be available to take its place, but with a reduced catalytic efficiency (91). On initial inspection, this might appear to be the case for Rep and UvrD, in light of the lower efficiency of block resolution for UvrD compared to Rep, noting that Rep deletion has a profound impact on replication whereas UvrD deletion does not (92). However, we find that Rep and UvrD do have different modes of colocation at a replication block. These differing mechanisms of replication promotion suggest a model in which different accessory helicases have evolved to fulfil the same overall function of resolving a block to DNA replication but in alternative ways. Even with different block removal efficiency, such redundancy may confer increased robustness, helping to ensure that DNA replication is carried out correctly.

We used single-molecule Slimfield to show that a high proportion of forks are colocated with UvrD *in vivo.* Although the raw imaging data is still subject to diffraction-limited resolution, localization tracking algorithms pinpoint fluorescently labelled UvrD to 40 nm lateral precision, smaller than the 50 nm diameter of the replisome machinery itself (61) and equivalent to ∼100 bp DNA. Thus, our measurements for UvrD and DnaQ are good indicators of genuine colocation of UvrD and the fork. Importantly, the degree of colocation we observed is much higher than expectations of random overlap of the optical images of DnaQ-mCherry and mGFP-UvrD.

Using SPR analysis we failed to find evidence for direct interaction partners at the replisome apart from the Tau protein. However, as a control, we tested Tau’s interaction with Rep and also found a positive interaction. Thus, we conclude that the interaction of UvrD with Tau is most likely non-specific and that UvrD recruitment to the replisome is mediated by blocks, rather than by a specific replisome interaction partner. A practical limitation of SPR measurement is that the immobilisation procedure does not normally preserve oligomer subunit interactions if they are not very tight, so one would expect to observe UvrD monomers on the surface. Where detection of an oligomer is specifically required, optimisation must be performed to stabilise it (e.g. cross-linking) for immobilisation. It is possible at very high immobilisation levels to have some “apparent” oligomer generated simply by the proximity of subunits but this is not a practically useful strategy. UvrD’s tetrameric form may also affect its interactions with replisome components. These limitations mean that SPR alone is semi-quantitative in not having the capability to report on the precise oligomeric state of UvrD on the SPR chip.

We probed UvrD’s function at the replication fork by perturbing its known DNA repair processing functions, knocking out *mutS* and *uvrA* to block mismatch and nucleotide excision repair respectively. We also introduced a mutant RNAP with *rpoB*35* and treated the wild type with rifampicin to reduce the occurrence of transcriptional blocks to translocating replication machinery. Deleting *uvrA* and rifampicin treatment produced clear results, reduced fork colocalisation, but deleting *mutS* and introducing the *rpoB*35* mutation did not. However, these latter perturbations did result in an identical phenotype of one extra UvrD tetramer per fork. Indeed, a recent study has confirmed the continuous physical association of UvrA and UvrD with RNAP. Our results are in agreement with their model proposed that UvrA and UvrD form a pre-TCRC that subsequently facilitates recruitment of UvrD at sites of DNA lesions.(50) The *rpoB*35* mutation improves cell viability to increased DNA lesions (83). Taken together, our results indicate that transcription-mediated lesions are a frequent impediment to active replisomes resulting in transcription-replication conflicts, that UvrD helps to clear via its association with RNAP. This intriguing link between mutations requires further study, possibly of these dual labelled strains in response to increased DNA lesions.

The only function of UvrD we did not probe directly was its role in RecA-mediated recruitment. We believe the results from a *recA* knockdown would be difficult to interpret due to its pleiotropic effects. RecA is the principal recombinase and induces the complex SOS response resulting in the induction of UvrD expression (93, 94). Furthermore, UvrD removes RecA from RecA-DNA complexes, thus functioning as an antirecombinase (95–97). Dissecting these UvrD functions from other effects that a *recA* mutation may have on the cell would be difficult. Applying the perturbations in combination, i.e. double knockdowns or knockdowns plus rifampicin, would likely lead to confusing results since too many deleterious perturbations may be lethal or change normal cell function too radically to be biologically relevant in the intended way. For the suite of mutant strains investigated we found small but significant colocalisation changes in UvrD, implying that UvrD-fork colocation is due to the frequent blocks to replication it encounters. Recent biochemical findings *in vitro* show that blocks to replication occur very frequently (1, 19, 98) potentially accounting for our observations here.

We show that forks stalled at transcriptional blocks can be rescued by addition of UvrD, prior to or post-stalling. If UvrD is to help perform this function then it must be present during the resolution of collision between the replication and transcription machinery, however, our estimate for dwell time at the fork of just a few milliseconds-compared to seconds for DnaQ (62)-suggests a non-negligible dissociation constant; UvrD is not processive *per se* but undergoes rapid turnover at the fork such that there is a high likelihood for UvrD being present at any given time, but not necessarily the same set of molecules. Interestingly, we observed a similar dwell time for Rep (13), despite it having a clear association partner in DnaB at the fork. However, using a defective restriction enzyme EcoRI E111G block produced a different effect, with late UvrD addition improving block resolution. This restriction enzyme block is artificial and less biologically relevant but may indicate that the known specific UvrD interaction with RNAP is important. Improved replication resolution with late addition of UvrD is intriguing, implying increased affinity for, or activity at, pre-stalled replication forks compared to UvrD being present as the fork stalls. Complete understanding requires further study of replication block resolution in different conditions.

Our findings suggest that UvrD diffuses and transiently interacts with DNA either at random, through interactions with RNAP, or interactions with a specific conformation of the blocked replication fork. At the DNA, UvrD is predominantly tetrameric, either pre-assembling in solution or on the DNA itself. The frequent sampling leads to UvrD often being in the vicinity of the replisome, allowing UvrD to resolve frequent blocks to replication. It is also possible that certain barriers to replication may be harder for Rep to resolve, and such barriers can be more efficiently removed by UvrD, however, it is still likely that they do not change the cell’s ability to replicate the genome in light of *uvrD* deletion still resulting in viable cells. Indeed, UvrD shares a high degree of structural similarity with Rep, although the two proteins differ significantly in their amino acid sequences (31, 99). While Rep is recruited to the fork through specific interactions with DnaB, no such interactions were found for UvrD. Stochastic impairment of the ability of Rep to interact with DnaB may necessitate block processing at the fork by UvrD. In summary, our results highlight the contribution of UvrD action to facilitating accurate genome duplication at active replisomes.

## Acknowledgments

We thank Maria Chechik for help with mGFP-UvrD purification. We thank Prof. Nigel Savery (University of Bristol) for the gift of the pETDuet-UvrD overexpression plasmid, and Dr Chris Hill (University of York) for the donation of ^32^P. We thank the Molecular Interactions Laboratory in the Bioscience Technology Facility at the University of York for technical assistance and support. Supported by BBSRC via grants BB/N006453/1 (M.L., P.M.), BB/R001235/1 (M.L.,P.M.), BB/N014863/1 (M.H.).

## Author Contributions

Conceptualization: AJMW, AS, PMG, MH, MCL

Methodology: AJMW, AS, JALH, AL, CG, PMG, MH, MCL

Software: AJMW

Investigation: AJMW, AS, JALH, APD, AL, DW, CG, MH

Visualization: AJMW, AS, MH

Supervision: AJMW, AS, MH, PMG, MCL

Writing—original draft: AJMW, AS, MH, MCL

Writing—review & editing: AJMW, AS, JALH, AL, DW, PMG, MH, MCL

## MATERIALS AND METHODS

### Strain construction

All strains used in this study (listed in full in SI Table S1, with associated plasmids and primers in SI Tables S2 and S3) are derivatives of the laboratory wild type strain TB28. Tagging of *dnaQ-mCherry-<kan>* (C-terminally labelled) is described in (13). *mGFP-uvrD-<kan>* (N-terminally labelled) was amplified from plasmid pAS79 (*uvrD^+^*) with primers oAS145 and oJGB383 having 50 bp homology to either end of the native *uvrD* locus. The resulting PCR product had homology either side such that recombination with the chromosome would result in integration of the PCR product at the native locus under the control of the native promoter. Prior to integration, all PCR products were treated with DpnI, gel purified, and introduced by electroporation into cells expressing the lambda Red genes from the plasmid pKD46 (55). The recombinants were selected for kanamycin resistance and screened for ampicillin sensitivity. The colonies obtained were verified for integration by PCR and sequencing. The *uvrD* recombinants were verified by PCR amplification using primers oPM319 and oPM320, and sequencing using the primers oJGB417, oJGB418, oPM407, oPM409, oPM411, and oMKG71. Where required, the kanamycin resistance gene was removed by expressing Flp recombinase from the plasmid pCP20 (100) to generate kanamycin sensitive strains carrying the fluorescent protein (FP) fusions. Dual labelled strains were created by introducing the kanamycin tagged FP alleles by standard P1 mediated transduction into single labelled strains carrying the required FP allele after removing the linked kanamycin marker.

### Determination of cell doubling time

*E. coli* strains were grown overnight in LB medium at 37°C at 200 rpm shaking. The saturated overnight cultures were diluted 100-fold into fresh LB or washed once with 1X 56 salts and diluted 100-fold in fresh 1X 56 salts with 0.2% glucose as the carbon source. Aliquots of 100 μl each of the diluted cultures in fresh media were pipetted into individual wells of 96 well clear flat bottom sterile microplates (Corning). The microplates containing the diluted cultures were incubated in a BMG LABTECH SPECTROstar Nano microplate reader at 37°C and the optical density (OD_600_) values were recorded at defined time intervals. The time taken for the optical density values to increase 2-fold during the exponential growth phase of the culture was taken as the cell doubling time (SI Table S4, Supplementary Figure S1).

### Protein Overexpression and Purification

*mGFP-uvrD* was excised from pAS79 using XhoI and SalI and ligated into pET14b cut with XhoI, creating pAS152 encoding N-terminal histidine-tagged mGFP-UvrD. pAS152 was used to overexpress the His-mGFP-UvrD fusion in BL21(DE3)/pLysS. A single colony was used to inoculate 10 mL of LB containing carbenicillin and chloramphenicol at 50 µg/mL and was incubated overnight at 37°C and 180 rpm. The overnight culture was clarified by centrifugation at 3,220 xg and 4°C, before the pellet was resuspended in 1 mL of LB and used to inoculate 1 L of LB containing carbenicillin and chloramphenicol at 50 µg/mL. The culture was then incubated at 37°C and 180 rpm until an OD_600_ of 0.6, at which point it was cooled to 20°C and IPTG was added to a 1 mM final concentration and incubation was continued at 20°C and 180 rpm overnight. Cells were pelleted by centrifugation at 3,500 xg and 4°C for 20 minutes, the pellet was resuspended in 20 mL of 50 mm Tris-HCl, pH 7.5, at 4 °C, and 10% sucrose (w/v) before flash freezing with liquid nitrogen and storing at-80°C.

Cells were thawed on ice and Tris pH 8.3, EGTA, EDTA, DTT, NaCl and lysozyme were added to final concentrations of 50 mM, 0.5 mM, 5 mM, 1 mM, 200 mM and 0.2 mg/mL respectively. The suspension was incubated on ice for 30 minutes before the addition of sodium deoxycholate to a final concentration of 0.05% (w/v) and allowed to stand on ice for a further 30 minutes. NaCl was then added to a final concentration of 500 mM before lysis by sonication on ice. The sample was clarified by centrifugation at 30,600 xg and 4°C for 20 minutes and the supernatant collected. The supernatant was subjected to polymin P precipitation, polymin P was added dropwise to a final concentration of 0.075% (w/v) under stirring at 4°C, stirring was continued for 10 minutes before centrifugation at 30,600g and 4°C for 20 minutes. The supernatant was collected and ammonium sulfate was added gradually to it whilst stirring at 4°C to a 50% saturation, the pellet was collected by centrifugation as previously and resuspended in binding buffer (20 mM Tris pH 8.3, 500 mM NaCl) + 5 mM imidazole. The sample was loaded onto a 5 mL HisTrap FF column that had been pre-equilibrated with binding buffer + 5 mM imidazole. The column was then washed with 15 column volumes of binding buffer + 5 mM imidazole before it was developed with a 20-column volume gradient of binding buffer + 5 to 300 mM imidazole. Peak fractions were monitored by their green colour as well as by SDS-PAGE, the peak fractions containing his-mGFP-UvrD were pooled and diluted in buffer A (20 mM Tris pH8.3, 1mM EDTA, 0.5 mM EGTA, 1mM DTT) to a conductivity equivalent to that of buffer A + 100 mM NaCl. The peak fractions were then loaded onto a 5 mL HiTrap Q column equilibrated with buffer A + 100 mM NaCl, the column was washed with 10 column volumes of buffer A + 100 mM NaCl and a gradient from 10 mM to 650 mM NaCl was developed over 20 column volumes. Peak fractions were assessed by SDS-PAGE and those containing his-mGFP-UvrD were loaded onto a HiLoad 26/60 Superdex 200 gel filtration column equilibrated with buffer B (20 mM Tris pH8.3, 300 mM NaCl), the column was developed with buffer B and peak fractions containing pure his-mGFP-UvrD as determined by SDS-PAGE were pooled and loaded onto a 5 mL HisTrap FF column that had been pre-equilibrated with binding buffer + 5 mM imidazole. The column was then washed with 3-column volumes of binding buffer + 5 mM imidazole before it was developed with a 6-column volume step of binding buffer + 200 mM imidazole. Peak fractions were then dialysed against storage buffer (20 mM Tris pH8.4, 300 mM NaCl, 10% glycerol) aliquoted and flash frozen in liquid nitrogen before storing at-80°C. Protein concentration was determined by Bradford’s assay.

UvrD was purified as previously described (56). Briefly UvrD was overexpressed in BL21-AI from a pETDuet vector, via induction by IPTG and arabinose. UvrD was purified from the soluble cell extract after lysis through the use of affinity (heparin agarose), anion exchange (Q-sepharose), and gel filtration chromatography. Pure UvrD was then stored at-80°C after dialysis into storage buffer or used immediately in Size Exclusion Chromatography-Multi-Angle Laser Light Scattering (SEC-MALLS) analysis or subsequently diluted to 1 nM for single-molecule *in vitro* experimentation.

Biotinylated UvrD and Rep were purified as previously described (12). Briefly, the proteins were overexpressed in a BL21 (DE3) containing pBirAcm, encoding a biotin ligase via induction by IPTG and the addition of biotin to the growth media. The biotinylated proteins were then purified from the soluble cell extracts by ammonium sulphate precipitation, and affinity chromatography (softlink avidin and heparin agarose). Pure proteins were then dialysed into storage buffer overnight and stored at-80°C

### Helicase assay

Assays for unwinding of streptavidin-bound forks were performed using a substrate made by annealing oligonucleotides oPM187B20 (5’ end labelled with ^32^P) and oPM188B34. Reactions were performed in 10 µL volumes containing 40 mM HEPES (pH 8); 10 mM DTT; 10 mM magnesium acetate; 2 mM ATP; 0.1 mg ml^-1^ BSA and 1 nM forked DNA substrate. Reactions were carried out as described earlier (13, 101). Briefly, the reaction mixture was pre-incubated at 37°C for five minutes with or without 1 µM streptavidin (Sigma-Aldrich), to which helicase (as indicated) and biotin (Sigma-Aldrich) were added to 100 µM (acting as a trap for free streptavidin) and incubated at 37°C for another 10 minutes. Reactions were stopped by addition of 2.5 µl of 2.5% SDS, 200 mM EDTA and 10 mg ml^-1^ of proteinase K. Reactions were then analysed by non-denaturing gel electrophoresis on 10% polyacrylamide TBE gels. The quantification of the unwinding and displacement of streptavidin from the fork was carried out as described in (102).

### Preparation of cells for microscopy

Cells were grown overnight in LB to saturation. The overnight cultures were inoculated the next day at 1:1000 dilution in 1X 56 salts supplemented with 0.2% glucose as the carbon source. The dilutions were then grown at 30°C to mid-log phase.

### Rifampicin treatment

Cells prepared for microscopy were treated with rifampicin at a final concentration of 50 µg/mL for 30 minutes at 30°C.

### Microscopy and image analysis

Imaging was performed on a bespoke dual-colour single-molecule microscope (13). Excitation from Obis 488 nm and 561 nm wavelength 50mW lasers (run at 20 mW) was reduced to 10 µm at full width half maximum field in the sample plane producing Slimfield (59) producing a mean excitation intensity of ∼0.25 mW/µm^2^. Lasers were digitally modulated to 5 ms period to produce alternating laser excitation using National Instruments dynamic I/O module NI 9402. Excitation was coupled into a Zeiss microscope body with the sample mounted on a Mad City Labs nanostage. Images were magnified to 80nm/pixel and imaged using an Andor Ixon 128 emCCD camera. Color channels were split red/green using a bespoke color splitter consisting of a dual-pass green/red dichroic mirror centered at long-pass wavelength 560 nm and emission filters with 25 nm bandwidths centered at 542 nm and 594 nm.

Cells were imaged on agarose pads suffused with minimal media as described previously (103). Foci were detected and tracked using previously described bespoke MATLAB software (104). In brief, bright candidate foci were detected in each frame by image transformation and thresholding based on Otsu’s method. A 17×17 pixel region of interest (ROI) is drawn around each candidate to define the local background, while a radius 5 circle is drawn in the center (the foreground) and subjected to iterative Gaussian masking (69). Foci were accepted if their signal to noise ratio was above 0.4, defined as the mean intensity inside the foreground divided by the standard deviation of the local background. Foci in successive frames were linked together into trajectories based on nearest distance, provided it was < 5 pixels. Linked foci were accepted as “tracks” nominally if they persist for at least 4 consecutive image frames.

Characteristic intensity of mGFP or mCherry was determined from the distribution of foci intensity values towards the end of the photobleach confirmed by overtracking foci beyond their bleaching and applying an edge-preserving filter (105, 106) to the raw intensity data to generate individual steps of the characteristic intensity, due to photobleaching (Supplementary Figure S7). This intensity was used to determine the stoichiometry of foci by fitting the first 3 intensity points with a straight line and dividing the intercept by this characteristic intensity. The number of peaks in the Gaussian fits to UvrD was set by running a peak fitting algorithm over the wild type distribution. This number of Gaussians was then used for mutant distributions unless two or more of the Gaussians converged on the same/similar peak value, in which case they were removed. For DnaQ, two peaks were fit as used previously (61). The width of the obtained Gaussian fits was in line with previous measurements (61) and a function of several sources of noise in the challenging live-cell environment, including: fluorophores broad intensity distribution (73) (Supplementary Figure S6); camera noise; cell autofluorescence; and mobility of the protein complexes themselves.

GFP and mCherry images were aligned based on the peak of the 2D cross correlation between their respective brightfield images. Colocalisation between foci and the probability of random colocalisation was determined as described previously (65, 107).

The microscopic apparent diffusion coefficients (*D*) were determined by fitting a straight line to the first three mean squared displacements (MSD) values constrained through the equivalent localization precision MSD as determined from the intensity (73, 108). *D* distributions were fit by three Gamma distributions as described previously (109–111). Dwell time was calculated as the number of frames that each trajectory was colocalised with the fork position, as determined by the DnaQ foci detected at time zero.

### Single-molecule *in vitro* imaging

10 µg/mL mouse anti-GFP (Roche. Cat no. 11814460001) in PBS was incubated on a tunnel slide for 5 min at room temperature, before washing with 100 µL of imaging buffer (20 mM Tris pH 7.8, 5 mM MgCl_2_, 1 mM DTT). 1 nM of purified mGFP-UvrD in imaging buffer was then flowed onto the slide and incubated at room temperature for 5 min before washing with 100 µL of imaging buffer. The ends of the channel were then sealed for imaging. We used a similar single-molecule fluorescence microscope design (107) which enabled Slimfield as for the live cell imaging in order to image the antibody immobilised mGFP-UvrD.

### SEC-MALLS

Size Exclusion Chromatography – Multi-Angle Laser Light Scattering (SEC-MALLS) experiments used Wyatt HELEOS-II multi-angle light scattering and rEX refractive index detectors linked to a Shimadzu HPLC system (SPD-20A UV detector, LC20-AD isocratic pump system, DGU-20A3 degasser and SIL-20A autosampler) with a G.E. Superdex S200 10/300 column at a flow rate of 0.5 mL/min in buffer of 20 mM Tris pH 8.3 (at 4degC) 1 mM EDTA, 1 mm EGTA, 1 mM DTT, 200 mM NaCl, 5% glycerol buffer. Data were analysed using Wyatt Astra 7 software. MWs were estimated using the Zimm fit method with degree 1. A value of 0.182 mL/g was used for protein refractive index increment (dn/dc).

### Surface plasmon resonance (SPR)

SPR was performed at 25°C on a BIAcore T200 instrument as in (12). Immobilisation of *E. coli* UvrD and Rep was performed onto streptavidin-coated SA sensor chips whilst the indicated concentrations of Tau and UvrB were passed over in 10 mM HEPES pH 7.4, 3 mM EDTA, 150 mM NaCl, 10 mM MgCl_2_ and 0.05% Tween 20 at 20 µl min^-1^. This buffer differed from that used in the DNA replication assays to minimise non-specific interactions with the surface-immobilised streptavidin. We performed experiments comparable to our previous study (12) in the range ca. 200-4000 nM.

### Replication assay

Replication assays were carried out using pPM872 as the template for RNAP blocks (1) or pPM594 (12) for EcoRI E111G blocks. Plasmid pPM872 contains P_lacUV5 52C_, a strong promoter in which the first 52 nucleotides of the transcript lack cytosine residues and are then followed by four consecutive cytosines. This enables transcription of P_lacUV5 52C_ to be stalled by the omission of CTP. Assays were performed in 40 mM HEPES (pH 8); 10 mM DTT; 10 mM magnesium acetate; 2 mM ATP; 0.2 mM GTP, 0.2 mM UTP; 0.04 mM dNTPs; and 0.1 mg/ml BSA. Reactions (15 µl) contained 2 nM plasmid template, 50 nM DNA polymerase III αɛθ complex, 25 nM τ clamp loader complex, 160 nM DnaB and DnaC monomers, 1 µM SSB, 80 nM β clamp, 30 nM HU, 200 nM DnaG, 300 nM DnaA. Helicases were added as indicated. Rep and UvrD used at 200 nM. *E. coli* RNAP holoenzyme from NEB (1 U/µl) was used at 1/3 dilution. Dilution was determined empirically to match RNAP replication inhibition levels from (1). EcoRI E111G was purified as in (112) and used at 200 nM. Reactions were assembled on ice and initiated by addition of DnaA and incubation for 4 min at 37°C, followed by addition of 60 units SmaI (Promega) plus 0.4 MBq [α^32^P] dCTP (222 TBq/mmol). Standard reactions with the helicase present in the initial protein mix were carried out at 37°C for 1 minute and then terminated by addition of 1 µl of 0.5 M EDTA. Delayed helicase addition was carried out by adding Rep/UvrD after 1 minute and incubation for a further minute at 37°C before termination with EDTA. Ethanol precipitated replication products were analysed by denaturing agarose gel electrophoresis (0.7% agarose in 2 mM EDTA, 30 mM NaOH for 400 volt hours, standard run was 16 hours at 25 V), phosphorimaging and autoradiography. 5’-labelled HindIII-digested lambda DNA was used as a marker. Gels were quantified using Quantity One® (Bio-Rad) software. Full-length plasmid replication products were quantified as a proportion of summed blocked products and normalised for length (for EcoRI E111G block), or total lane signal (RNAP block).

## Supplementary Information

## Supplementary Tables

**Table S1.** **Strains used in this study**

| Strain | Relevant genotype | Source or derivation |
| --- | --- | --- |
| BW25113 derivatives: |  |  |
| BW25113 | *rrnB* Δ*lacZ*4787 *hsdR*514 $\Delta$(*araBAD*)567 $\Delta$(*rhaBAD*)568 *rph*-1 | (1) |
| JW2703 | Δ*mutS*::<kan> | (1) |
| JW3786 | Δ*uvrD*::<kan> | (1) |
| JW4019 | Δ*uvrA*::<kan> | (1) |
| MG1655 derivatives: |  |  |
| TB12 | Δ*lacI*ZYA::<kan> | |
| TB28 | Δ*lacI*ZYA | (2, 3) |
| TB28 derivatives |  |  |
| N6577 | Δ*lacI*ZYA Δ*rep*::*cat* | (4) |
| N6661 | Δ*lacI*ZYA::<> Δ*rep*::*dhfr* / pAM407 (*uvrD*<sup>+</sup> *lacZ*<sup>+</sup>, a pRC7 derivative) | (4) |
| N6661 | Δ*lacI*ZYA::<> Δ*rep*::*dhfr* / pAM407 (*uvrD*<sup>+</sup> *lacZ*<sup>+</sup>, a pRC7 derivative) | (4) |
| N6644 | Δ*lacI*ZYA::<> *uvrD*::*dhfr* Δ*rep*::*cat* / pAM407 (*uvrD*<sup>+</sup> *lacZ*<sup>+</sup>, a pRC7 derivative) | (4) |
| JGB166 | Δ*uvrD* / pKD46 | JW3786 transformed with pKD46 |
| AS97 | TB28/pKD46 | (5) |
| AS446 | *dnaQ*-*mCherry*-<kan> |  |
| AS448 | *dnaQ*-*mCherry*-<> | (5) |
| AS512 | *mGFP*-*uvrD*-<kan> | *mGFP*-*uvrD*-<kan> recombineered into JGB166 |
| AS517 | Δ*lacI*ZYA::<> Δ*rep*::*cat* *mGFP*-*uvrD*-<kan> / pAM407 (*uvrD*<sup>+</sup> *lacZ*<sup>+</sup>, a pRC7 derivative) | N6661 X P1(AS512) |
| AS564 | *dnaQ* - *mCherry* -<kan> *rpoB* *35 | PM486 X P1(AS446) |
| AS571 | *dnaQ* - *mCherry* -<> *rpoB* *35 | AS564 to km<sup>S</sup> with pCP20 |
| AS580 | *dnaQ*-*mCherry*-<> *mGFP*-*uvrD*-<kan> | AS448 X P1(AS512) |
| AS582 | *dnaQ* - *mCherry* -<> *mGFP* - *uvrD* -<kan> *rpoB* *35 | AS571 X P1(AS512) |
| AS590 | *dnaQ*-*mCherry*-<> *mGFP*-*uvrD*-<kan> Δ*rep*::*cat* | AS580 X P1(N6577) |
| AS650 | *dnaQ*-*mCherry*-<> *mGFP*-*uvrD*-<> | AS580 to km<sup>S</sup> with pCP20 |
| AS656 | *dnaQ*-*mCherry*-<> *mGFP*-*uvrD*-<> Δ*uvrA*::<kan> | AS650 X P1(JW4019) |
| AS658 | *dnaQ*-*mCherry*-<> *mGFP*-*uvrD*-<> Δ*mutS*::<kan> | AS650 X P1(JW2703) |

**Table S2.**
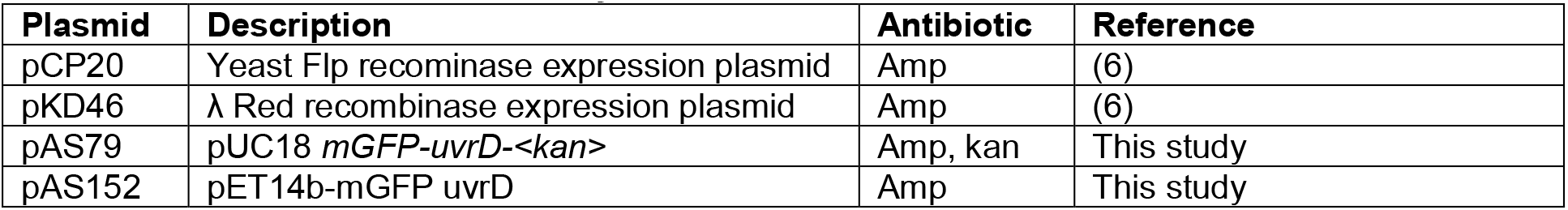
**Plasmids used in this study**

**Table S3.**
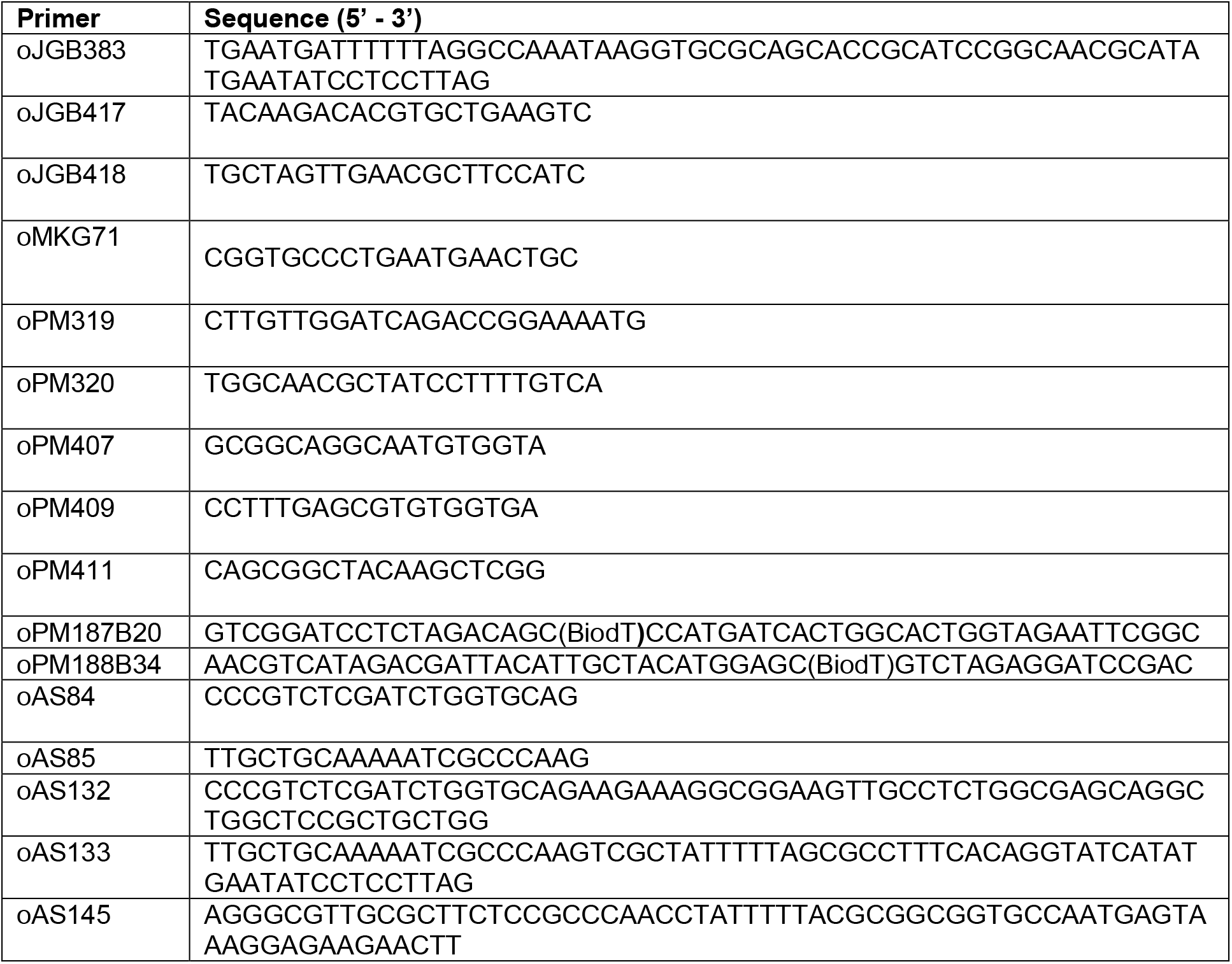
**Oligonucleotides used in this study**

**Table S4.** **Cell doubling times of labelled strains.** Values expressed are the means of three independent replicates, with standard deviation and standard errors indicated.

| Strain (Relevant genotype) | LB medium |  |  | Minimal medium |  |  |
| --- | --- | --- | --- | --- | --- | --- |
|  | Mean doubling time (min) | SD (min) | SEM (min) | Mean doubling time (min) | SD (min) | SEM (min) |
| WT | 23.72 | 1.11 | 0.64 | 62.80 | 17.06 | 9.85 |
| *dnaQ-mCherry* | 26.85 | 0.91 | 0.52 | 62.22 | 18.63 | 10.76 |
| *mGFP-uvrD* | 24.79 | 0.37 | 0.22 | 59.45 | 1.80 | 1.04 |
| *dnaQ-mCherry mGFP-uvrD* | 25.27 | 1.51 | 0.87 | 71.03 | 7.59 | 4.38 |
| $\Delta mutS$ | 24.59 | 0.55 | 0.32 | 107.91 | 13.96 | 8.06 |
| $\Delta uvrA$ | 27.99 | 0.23 | 0.13 | 92.25 | 5.86 | 3.38 |
| $\Delta rpoB^{*35}$ | 33.98 | 0.17 | 0.10 | 83.11 | 30.83 | 17.80 |
| $\Delta rep$ | 24.30 | 2.03 | 1.17 | 62.74 | 1.20 | 0.70 |
| *dnaQ-mCherry mGFP-uvrD $\Delta mutS$* | 26.85 | 0.68 | 0.39 | 110.84 | 34.91 | 20.16 |
| *dnaQ-mCherry mGFP-uvrD $\Delta uvrA$* | 24.13 | 0.56 | 0.33 | 91.80 | 5.18 | 2.99 |
| *dnaQ-mCherry mGFP-uvrD $\Delta rpoB^{*35}$* | 32.81 | 0.70 | 0.40 | 99.11 | 78.68 | 45.42 |
| *dnaQ-mCherry mGFP-uvrD $\Delta rep$* | 26.54 | 1.90 | 1.10 | 62.17 | 2.98 | 1.72 |

**Table S5.** **Comparison of wild type and labelled UvrD**

| Method | Figure | Result |
| --- | --- | --- |
| Growth rates | S1 | No significant difference |
| Plasmid loss assays | S2 | mGFP-UvrD can rescue cells where UvrD deletion is lethal |
| *In vitro* unwinding assay | S3 | Nucleoprotein block removal and helicase activity of mGFP-UvrD and biotinylated UvrD was comparable to the wild type |
| SEC-MALLS | S9 | No significant detectable difference in oligomeric state |
| *In vitro* replication assay | Fig. 5 and S14 | Wild type UvrD and mGFP-UvrD are able to overcome blocks to replication |

## Supplementary figures

**Supplementary Figure S1.**
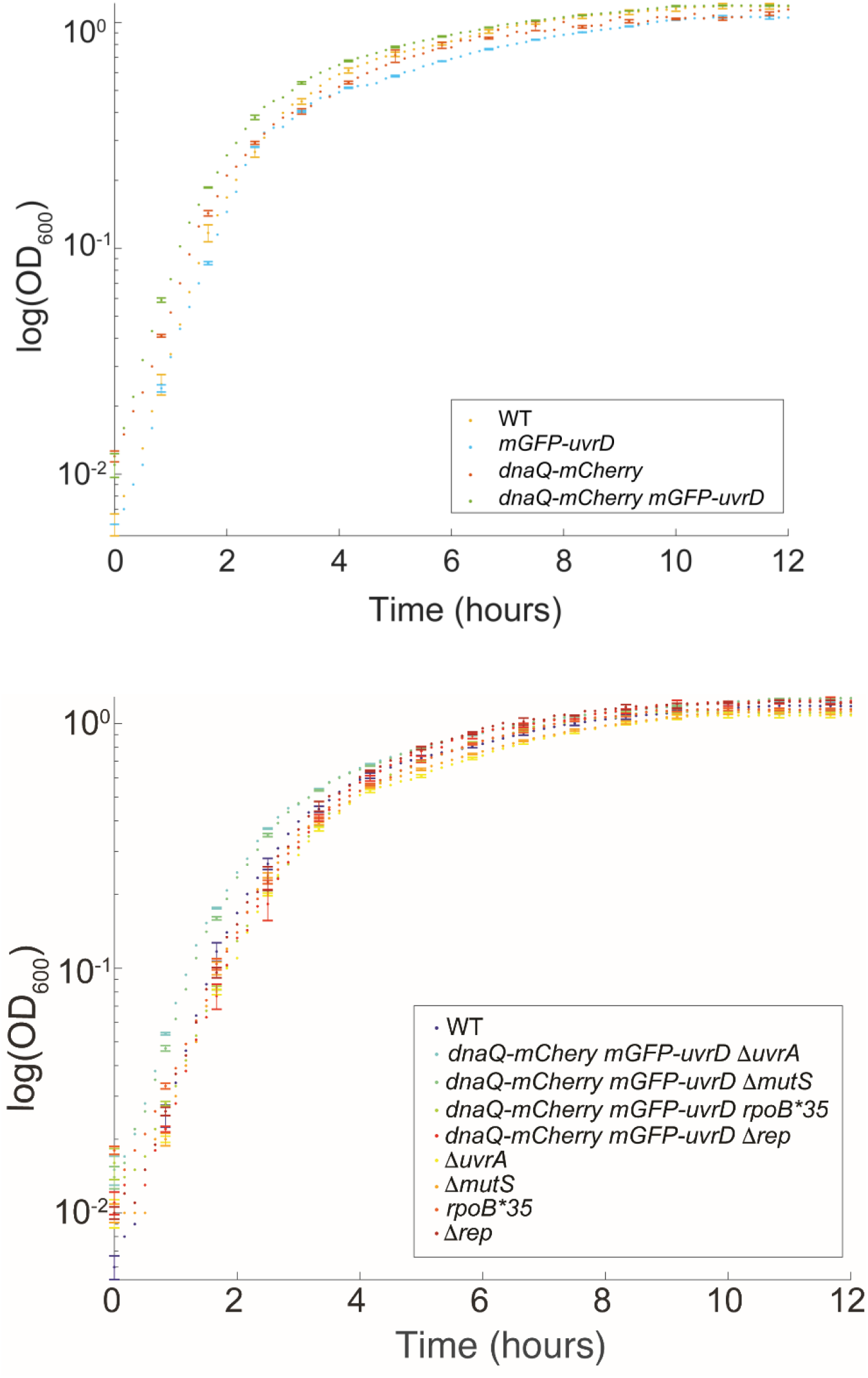
Growth curves in LB medium. Cultures were grown in LB as described in the text. The OD600 values are plotted on a log scale on the vertical axis with a linear scale of time in hours on the horizontal axis. The genotypes of the strains are indicated in boxes within the figures. SD error bars (shown just on every 5^th^ consecutive point for clarity here), taken from N=3 replicate cultures. Plots in panel A are growth curves of single and dual labelled wild-type strains, while those in panel B are dual labelled and unlabelled mutant strains. The unlabelled wild-type strain is included in all panels as a reference.

**Supplementary Figure S2.**
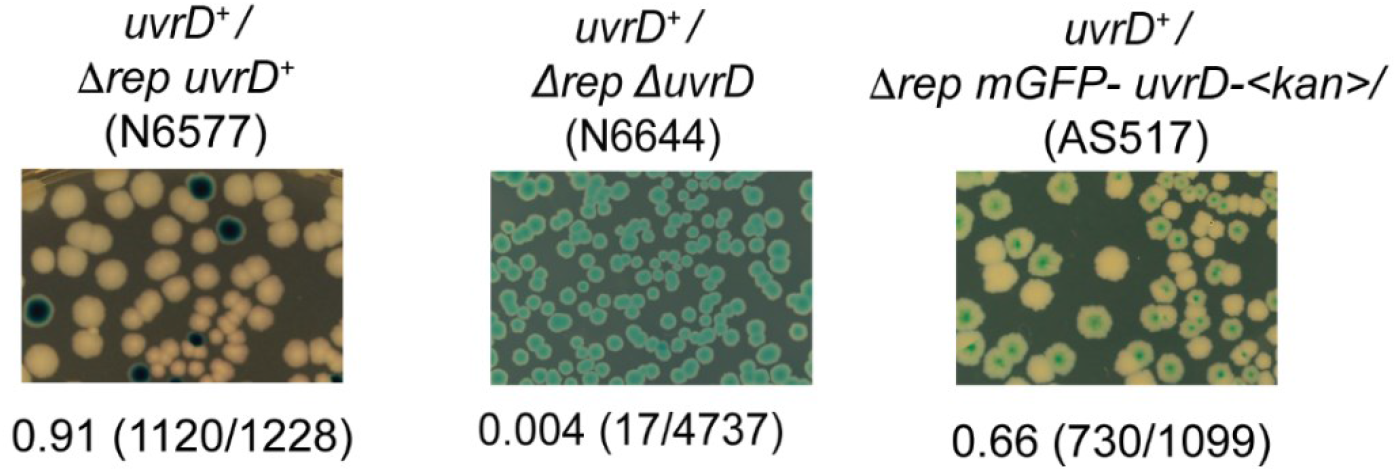
Testing of mGFP-UvrD fusions for retention of function. We transformed the strain carrying a chromosomal *mGFP-uvrD* allele with a derivative of pRC7 carrying a wild-type *uvrD* allele (pPM407), which is an unstable low copy plasmid that also carries the *lacZYA* genes (3). Presence of this plasmid confers blue colour to the colonies on plates containing Xgal when strains are chromosomally deleted for the *lac* operon. In rapidly growing cells, cells require either *rep* or *uvrD* for viability, and the loss of both is lethal (4, 7). Thus, *Δrep* cells with a functional *uvrD* allele are viable and can lose pRC7*uvrD*, evidenced by appearance of white colonies on LB Xgal IPTG plates, but cells lacking *uvrD* function cannot survive upon loss of the plasmid resulting in recovery of only blue colonies. The *Δrep mGFP-uvrD* cells produced white colonies on Xgal media, indicating that the *mGFP-uvrD* fusion is functional.

**Supplementary Figure S3.**
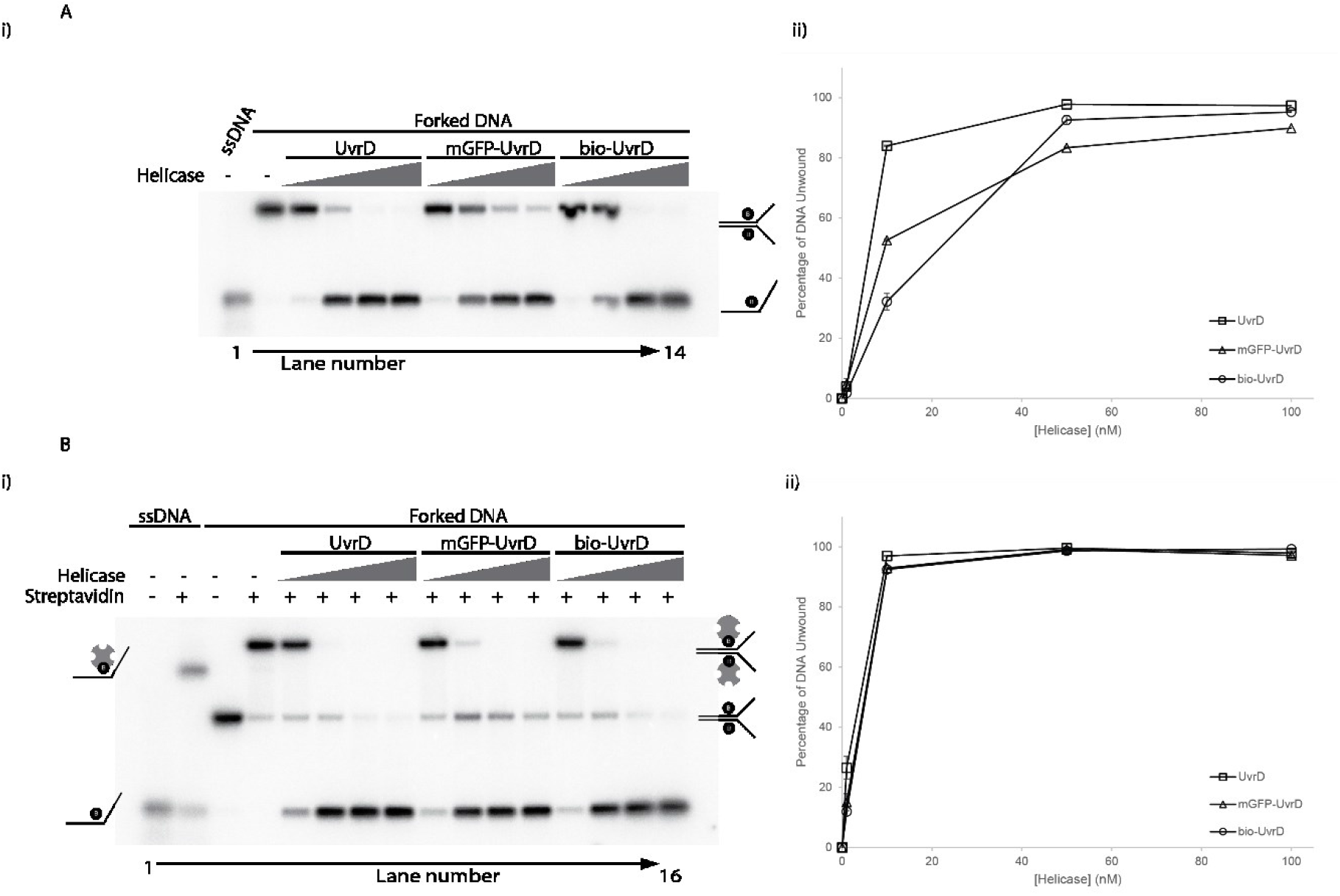
mGFP-UvrD fusion is functional *in vitro*. (A)(i) Representative native polyacrylamide TBE gel showing UvrD, mGFP-UvrD and biotinylated-UvrD unwinding of forked DNA containing biotin on both strands within the duplex region (8, 9). Lanes 1-2 contain markers indicating the position of single stranded or forked DNA as indicated. Lanes 3-14 contain the products of unwinding the forked DNA by the indicated helicase at 1, 10, 50 and 100 nM. (ii) Quantification of the unwinding of the forked substrate by the indicated helicases. Data points shown are averages of band intensities (taken from two gels) with standard errors indicated by bars on the points. (B)(i) Representative native polyacrylamide TBE gels showing UvrD, mGFP-UvrD and biotinylated-UvrD unwinding of forked DNA containing biotin on both strands within the duplex region (8, 9). Lanes 1-4 contain markers indicating the position of single stranded or forked DNA +/-streptavidin as indicated. Lanes 5-16 contain the products of unwinding the forked DNA +/-streptavidin by the indicated helicase at 1, 10, 50 and 100 nM. (ii) Quantification of the unwinding of the forked, streptavidin bound substrate by the indicated helicases. Data points shown are averages of band intensities (taken from two gels) with standard errors indicated by bars on the points.

**Supplementary Figure S4.**
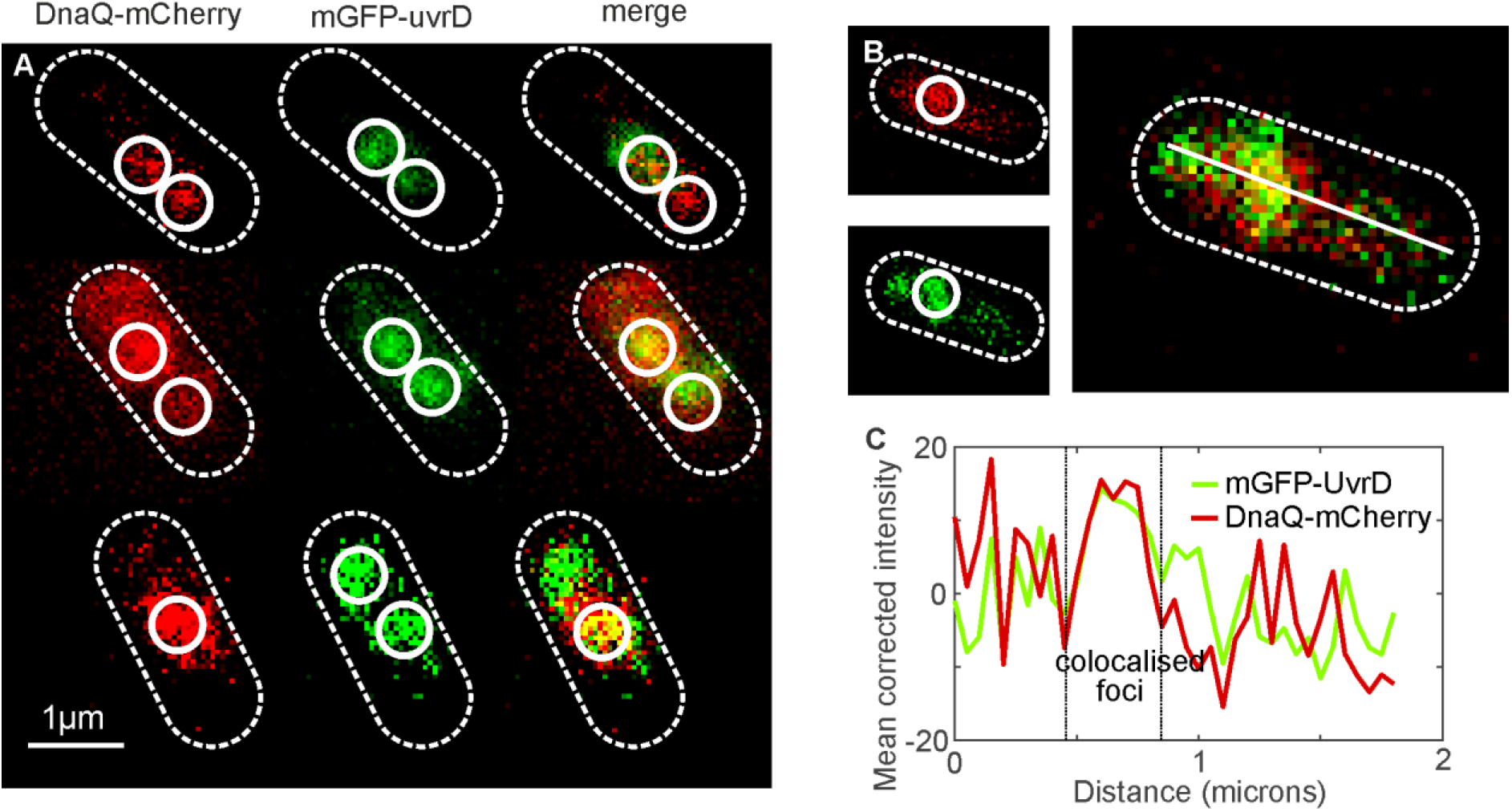
Cells typically appear to have two DnaQ foci, but more rarely show just a single DnaQ focus corresponding to two forks separated by less than the optical resolution limit. A. and B. Dual color Slimfield of mGFP-UvrD:DnaQ-mCherry, scale bar 1 micron. Segmented outlines of cell bodies (white dash) and detected foci (white circles) indicated. C. Mean corrected intensity line profile through a colocalized foci in B.

**Supplementary Figure S5.**
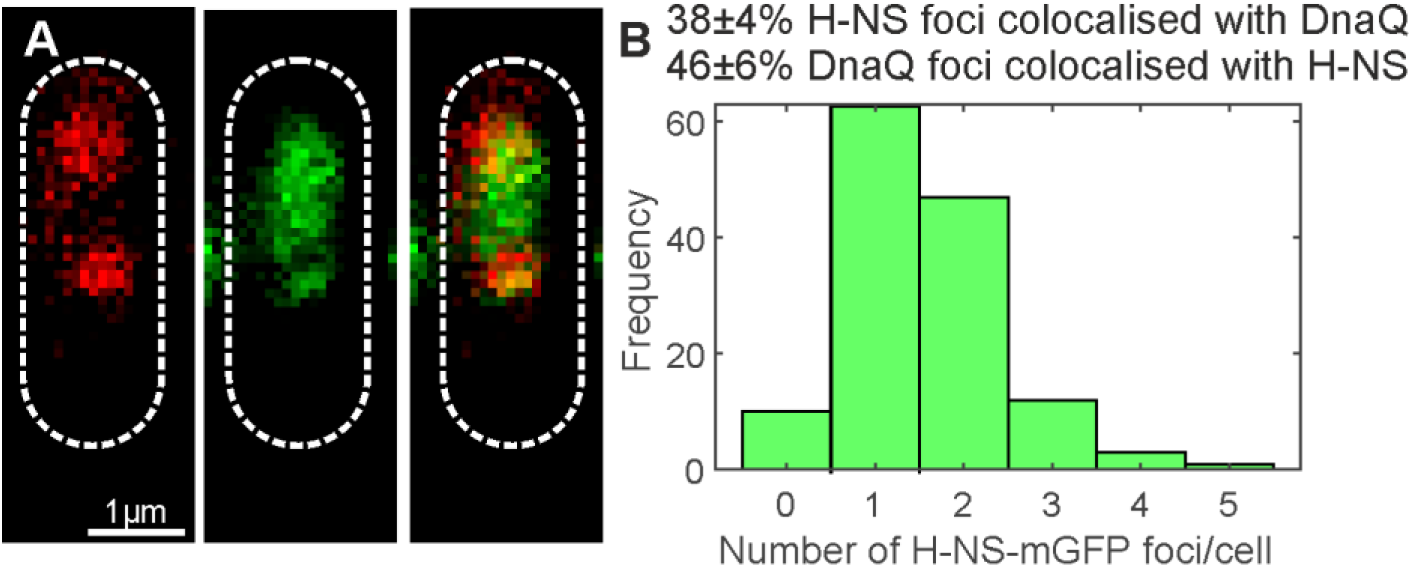
H-NS associates with the nucleoid but not specifically with the fork. A. Dual color Slimfield of H-NS-mGFP-:DnaQ-mCherry, scale bar 1 micron. Segmented outlines of cell bodies indicated (white dash). B. Histogram showing the number of H-NS foci detected per cell, SD errors.

**Supplementary Figure S6.**
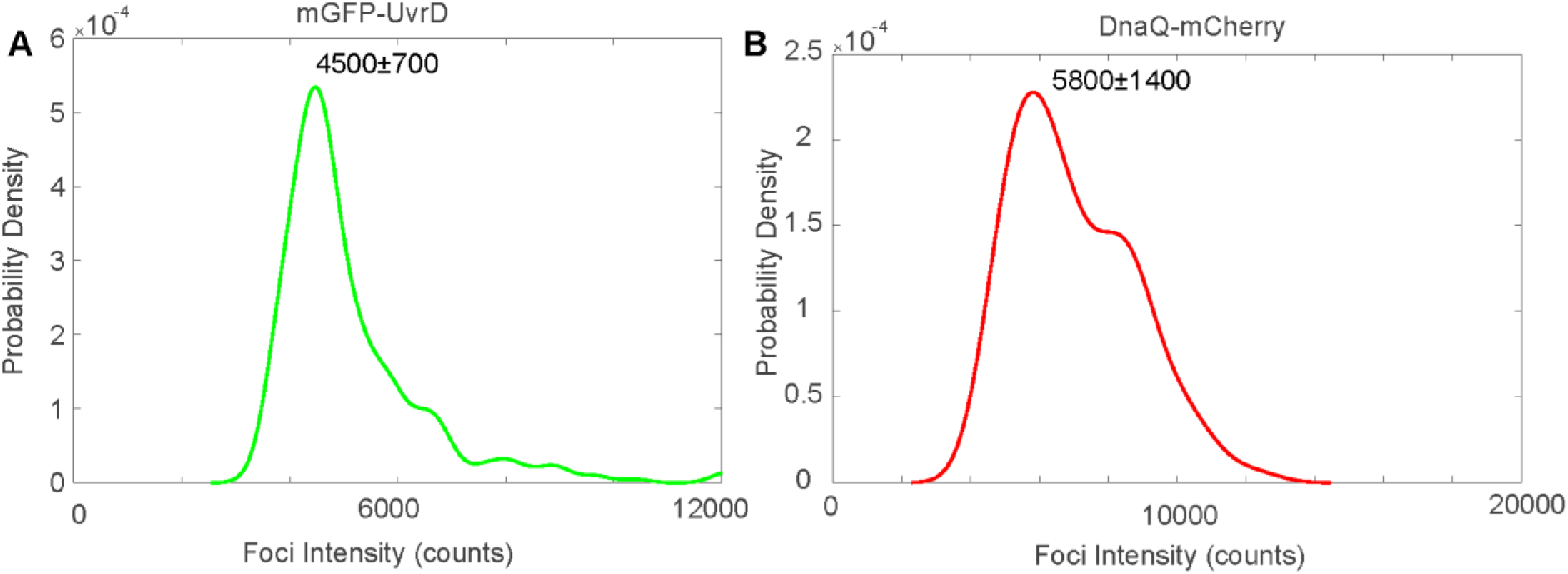
Brightness of single mGFP and mCherry molecules. Characteristic intensity distributions rendered as kernel density estimates of single A. mGFP-UvrD and B. DnaQ-mCherry obtained using Slimfield. Peak±full width at half maximum indicated. Distributions calculated from the tracked foci intensity distributions from the end of the photobleach process such that only single fluorophore molecules are detected. Number of molecules per foci before bleaching is determined by dividing the initial foci intensity by these values for the equivalent fluorophore.

**Supplementary Figure S7.**
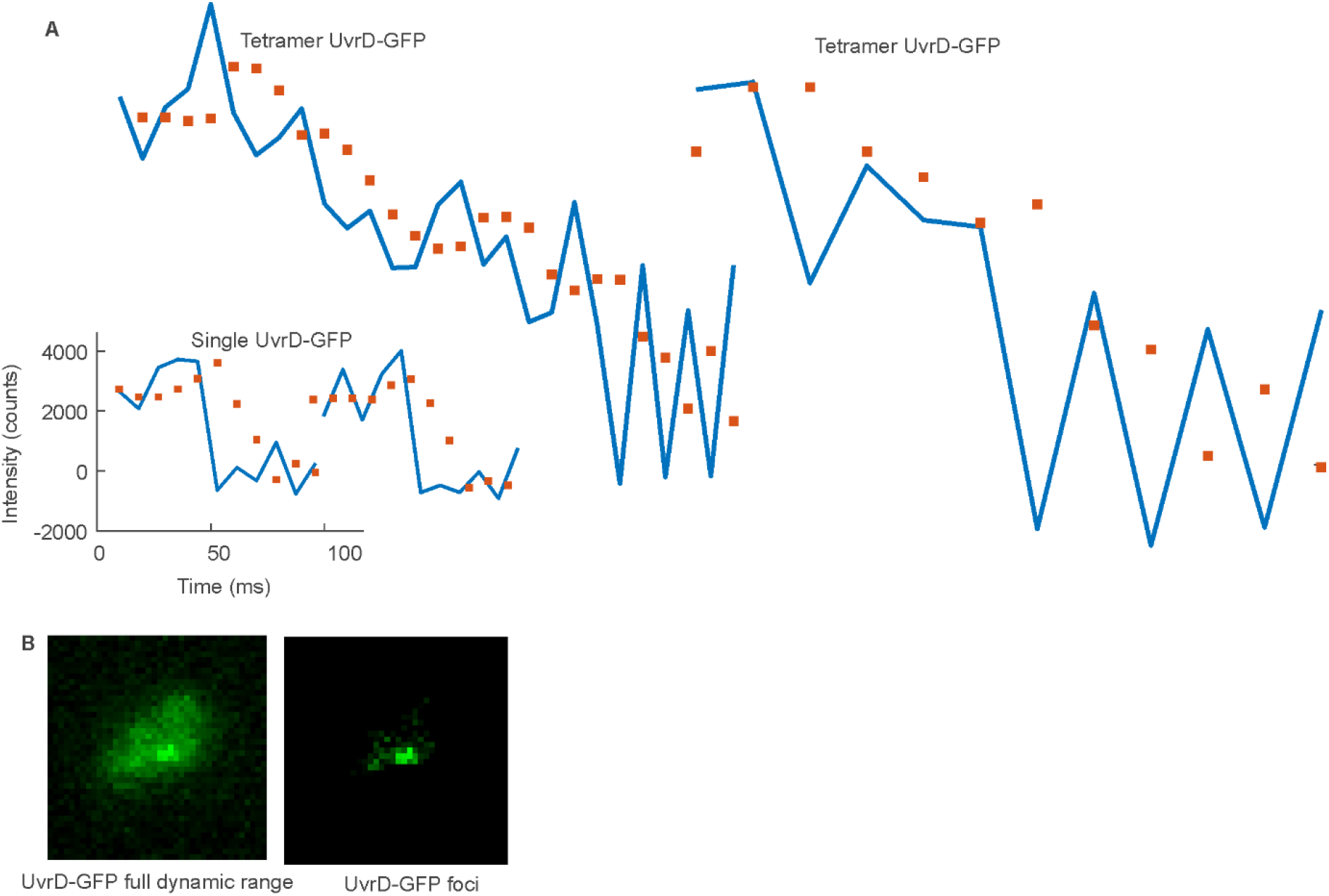
Single-molecule image analysis. **A.** Foci intensity as a function of time for detected UvrD tracks (blue) overlaid by Chung-Kennedy (CK) filtered intensity (squares). CK filtering is an edge preserving filter allowing noise to be filtered out while retaining steps.(113) Examples are shown of intensity traces for single UvrD tracks showing tetramers exhibiting multiple photobleach steps and monomers detected towards the end of photobleaching. Foci were ‘overtracked’ beyond the last detected bright frame to illustrate steps. **B**. UvrD-GFP micrograph showing the full dynamic range (left) and adjusted to illustrate a foci (right)

**Supplementary Figure S8.**
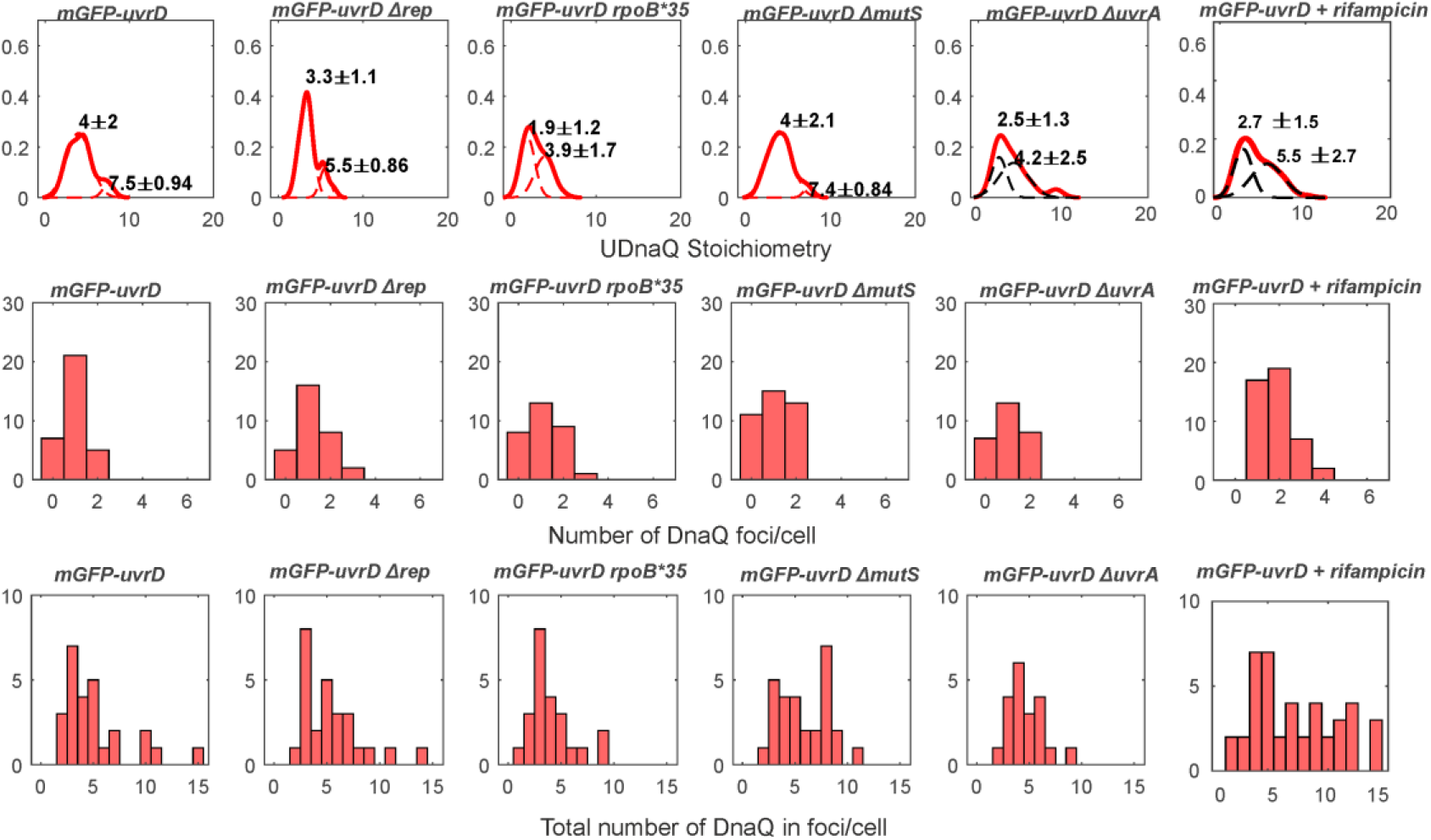
DnaQ tracking characterization. Top row: Distribution of DnaQ foci stoichiometry as a kernel density estimate for all strains and conditions (red) with two Gaussian fit overlaid as black dotted lines. Peak ± half width at half maximum shown above. Middle row: Corresponding histogram of number of foci detected per cell. Bottom row: histogram of the total amount of DnaQ in foci per cell. N=30-50 cells per strain or condition.

**Supplementary Figure S9.**
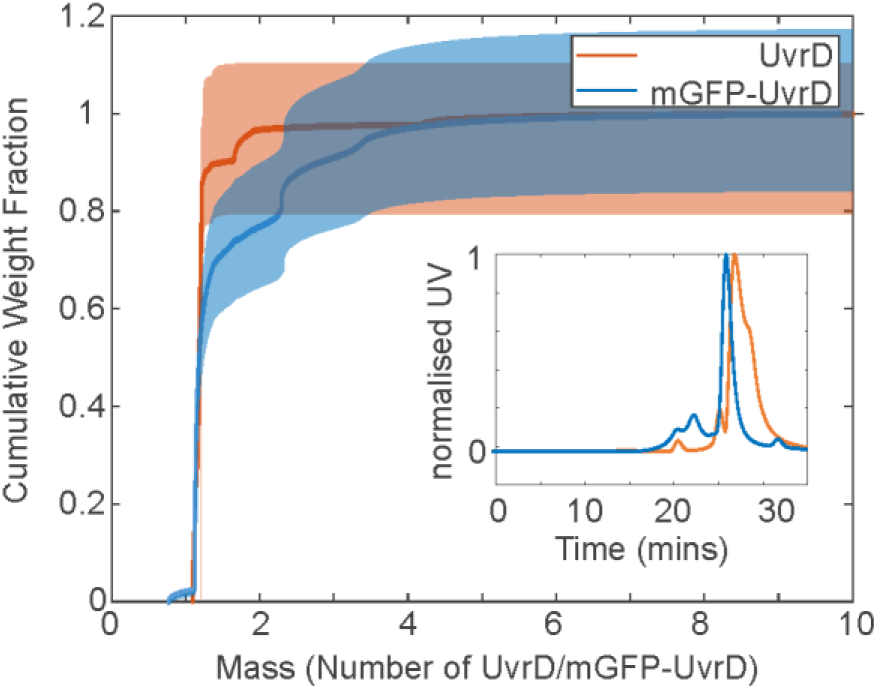
SEC-MALLS of UvrD/mGFP UvrD. Size Exclusion Chromatography-Multi-Angle Laser Light Scattering (SEC-MALLS) of 4 µM UvrD (red) and mGFP-UvrD (blue) cumulative weight fraction as a function of molar mass in number of UvrD/mGFP-UvrD, shading indicates 16% indicative error bounds consistent with sensitivity error measurements reported from polydispersed heterogenous samples using SEC-MALLS (68), and consistent with our own estimation of sensitivity using our SEC-MALLS system. Inset shows the normalised UV absorbance as a function of elution time through the column with the monomer peaks indicated.

**Supplementary Figure S9.**
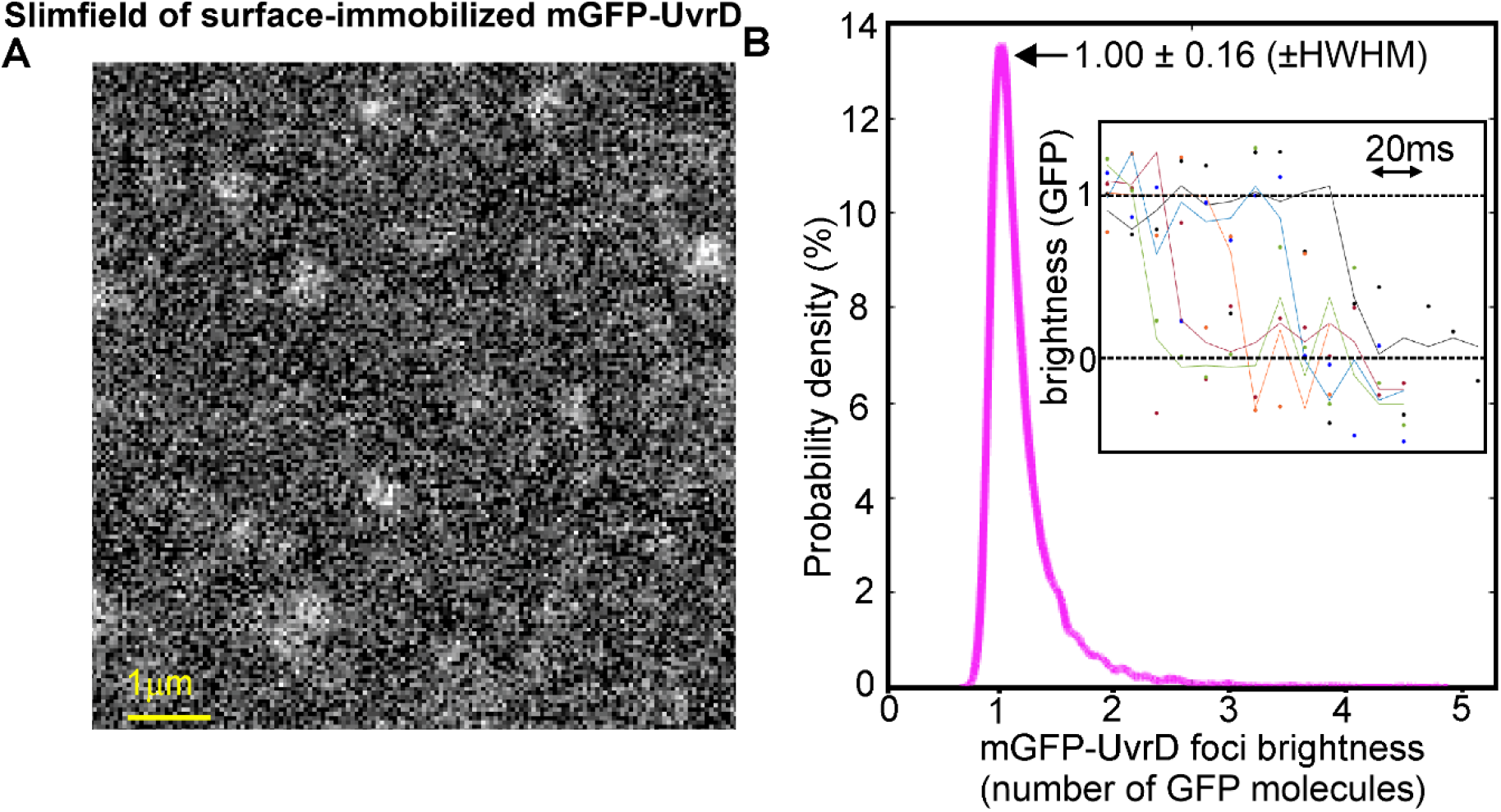
Single-molecule microscopy of purified mGFP-UvrD indicates most foci contain monomeric mGFP-UvrD. A. Representative Slimfield of surface-immobilized mGFP-UvrD. B. Automated single-particle tracking output for mGFP-UvrD showing probability distribution for mGFP-UvrD foci brightness normalized by the single-molecule dye brightness, peak (±HWHM) corresponding to 1 molecule indicated (arrow) for n=7,384 foci. *Inset*: example overlaid stepwise photobleach traces for four different mGFP-UvrD foci indicating a consistent single-molecule step to zero brightness whose size is comparable to the measured modal brightness across all foci (dots: raw data, lines: Chung-Kennedy edge-preserving filter output).

**Supplementary Figure S10.**
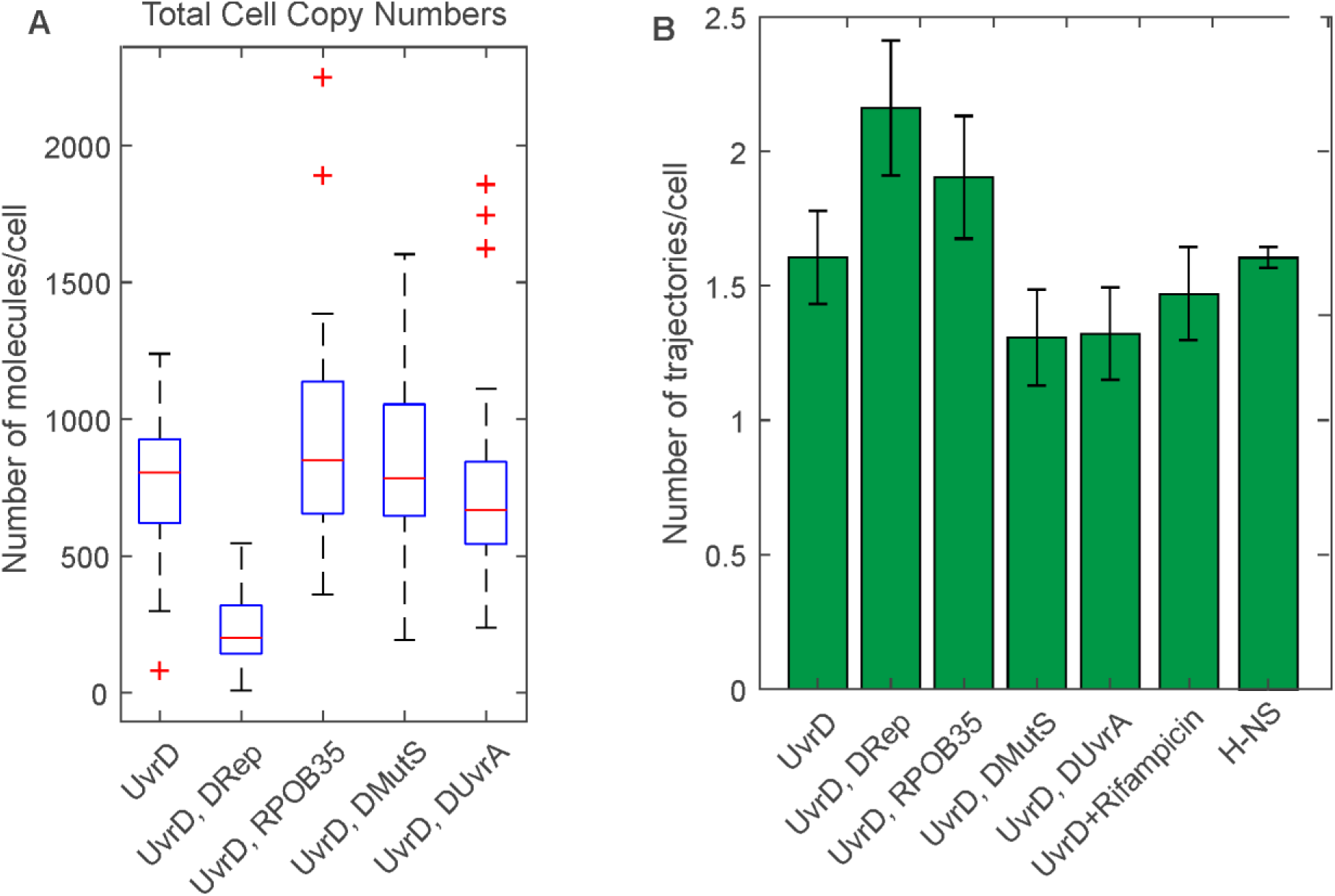
A. Boxplot of the total number of mGFP-UvrD molecules per cell, estimated by numerical integration of the whole cell fluorescence. Median is shown as red line, bottom and top of the blue box mark the 25^th^ and 75^th^ percentiles and whiskers extend to the most extreme points not considered outliers (2.7 standard deviations covering 99.3% of normally distributed data) with outliers beyond this shown as red +. N=30 cells per strain. B Number of trajectories per cell used in the stoichiometry determination

**Supplementary Figure S11.**
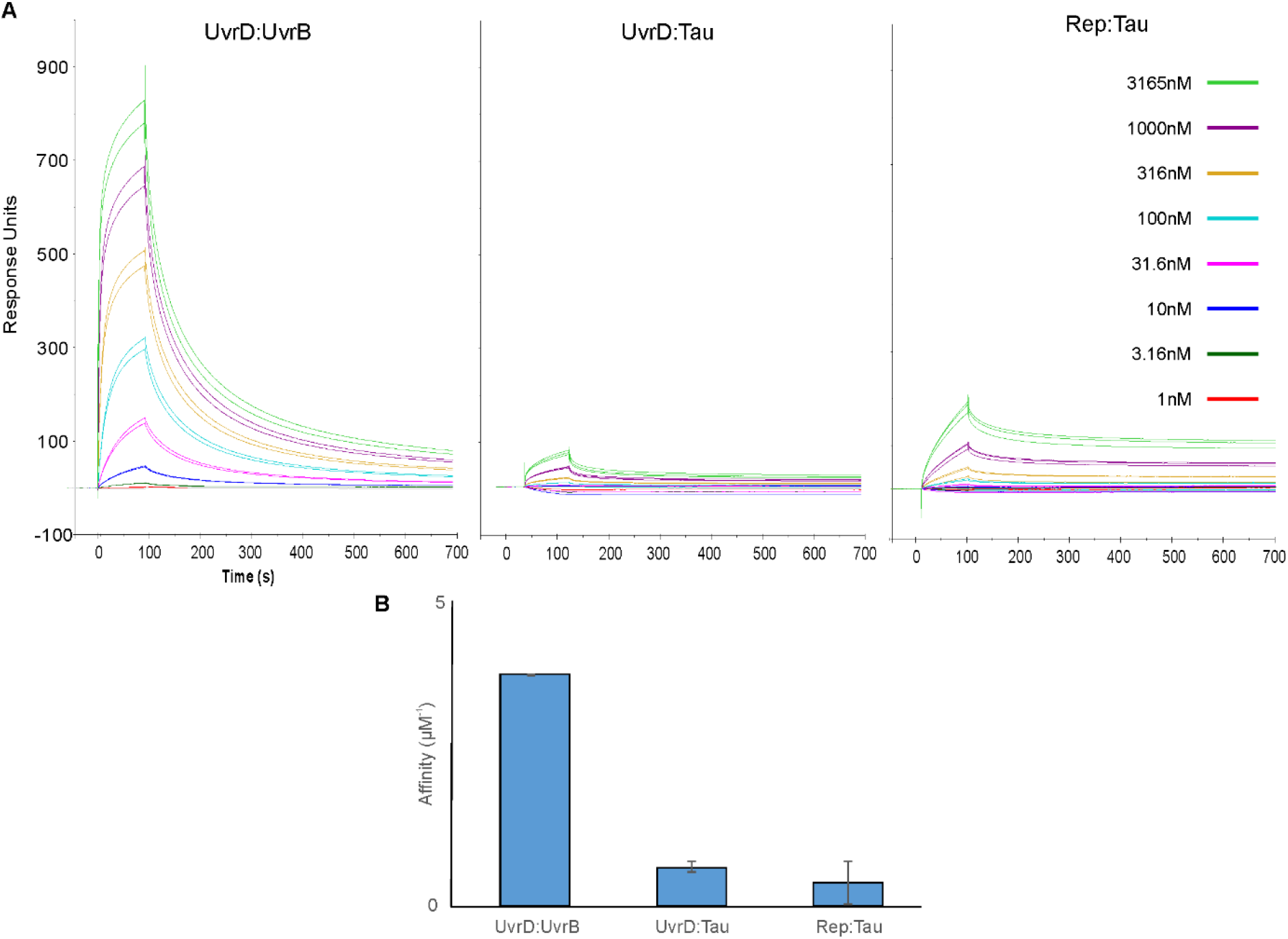
Surface plasmon resonance measurements of helicase interactions with Tau. A. Titration response sensorgrams for UvrD:UvrB, UvrD:Tau and Rep:Tau with indicated analyte concentrations. B. Binding affinity plotted as association constant (Ka, uM-1) for the UvrD:UvrB, UvrD:Tau and Rep:Tau interactions assuming a single sites binding model. The respective dissociation constants Kd are 231 nM, 1.4 µM and 2.3 µM.SD error bars.

**Supplementary Figure S12.**
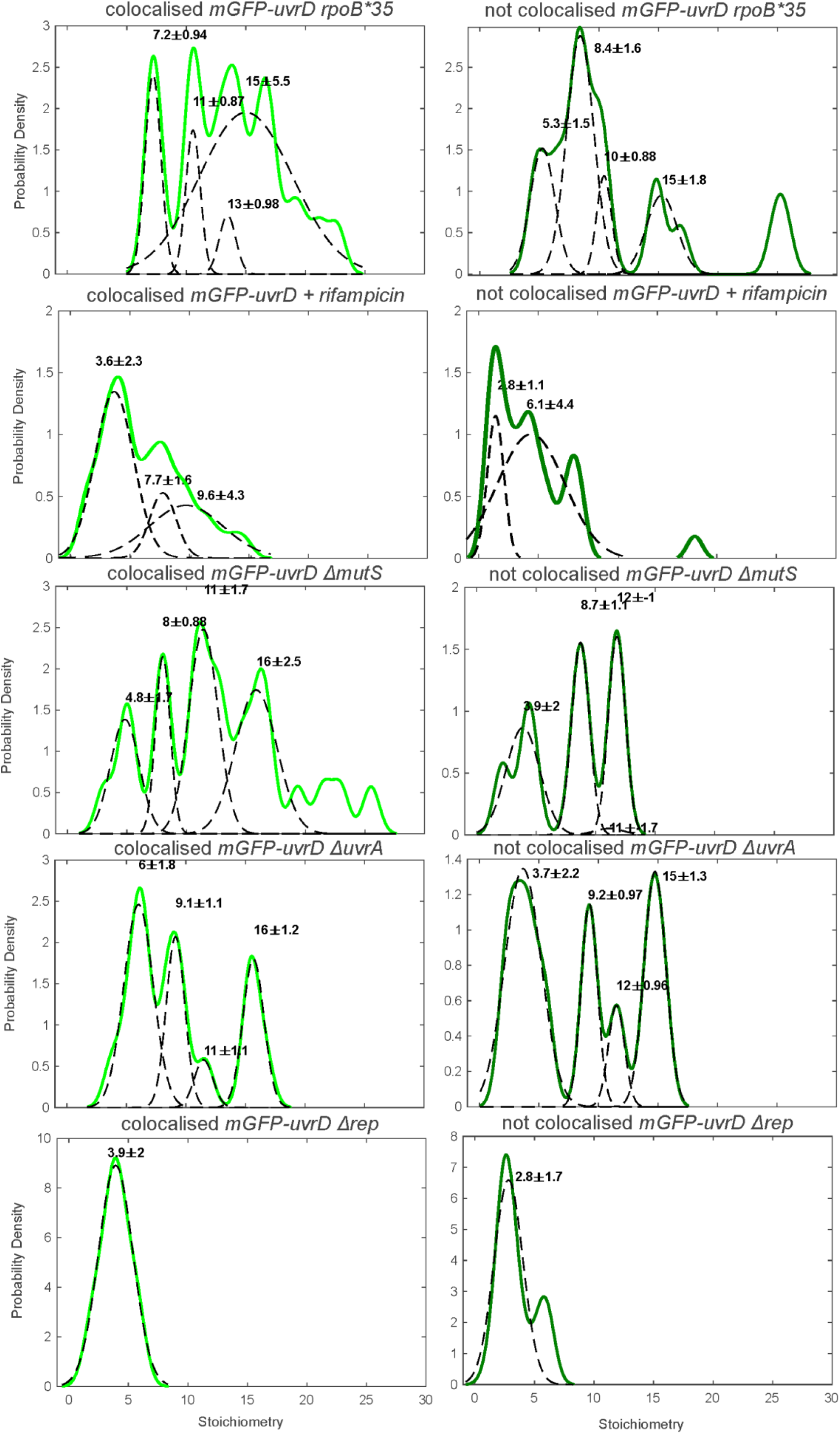
Perturbed UvrD stoichiometry. The distribution of UvrD foci stoichiometry rendered as a kernel density estimate for foci colocalised (Left) and non-colocalised (Right) with DnaQ foci. Distributions were fitted with multiple Gaussian fits (black dotted lines) with the peak value ± half width at half maximum indicated above each peak.

**Supplementary Figure S13.**
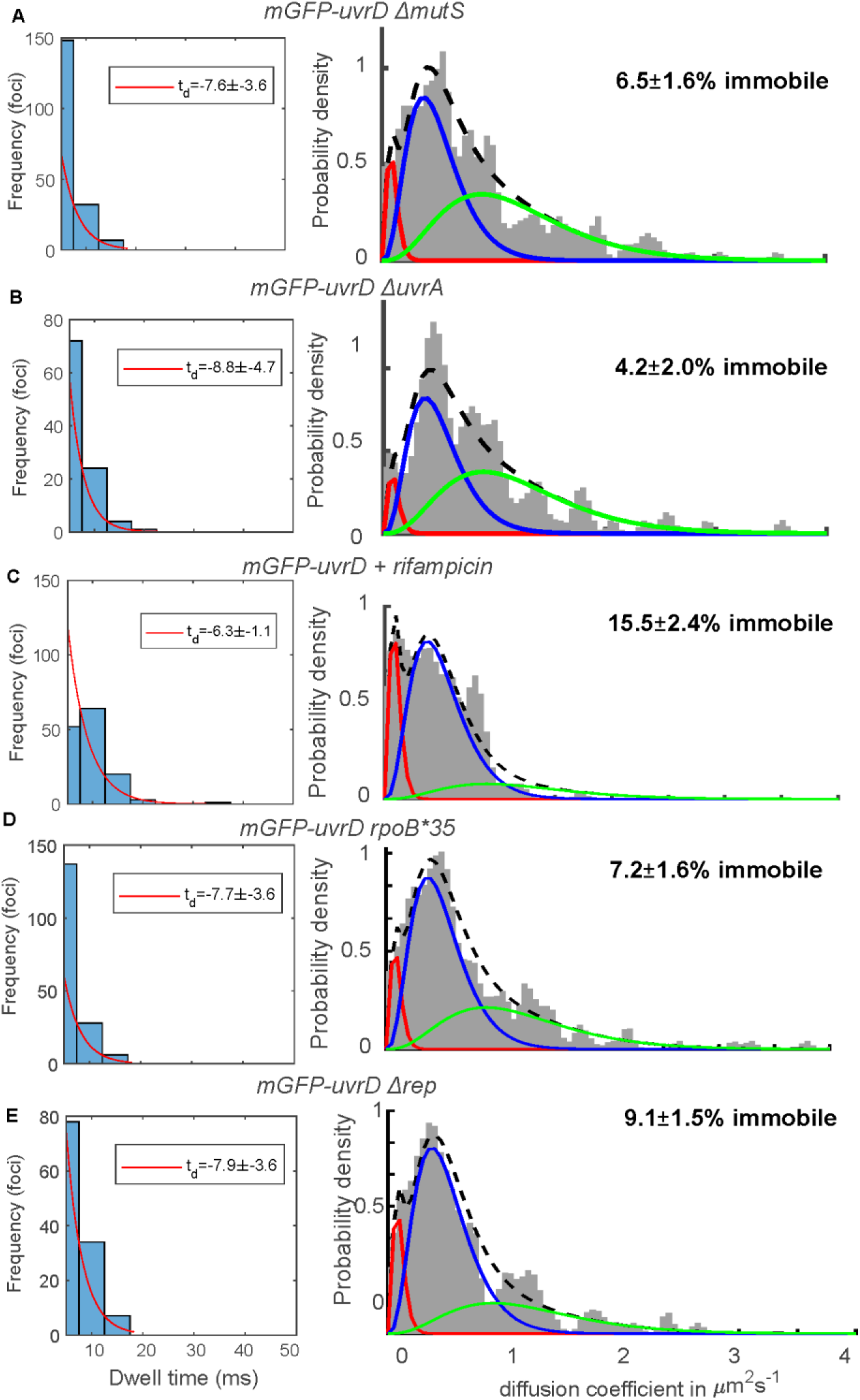
UvrD perturbed dynamics. A-E Left, distribution of time UvrD and DnaQ foci were colocalised (blue) fit with an exponential (red) to yield a characteristic dwell time. Right, distribution of microscopic diffusion coefficients of UvrD foci (grey) fitted with a three-state gamma distribution model containing a relatively immobile population with *D* = 0.1 µm^2^/s (red), a transiently immobile population, *D* = 0.5 µm^2^/s (blue) and a mobile population, *D*=1.2 µm^2^/s (green). Errors represent 95% confidence intervals. N∼300 foci/condition.

**Supplementary Figure S14.**
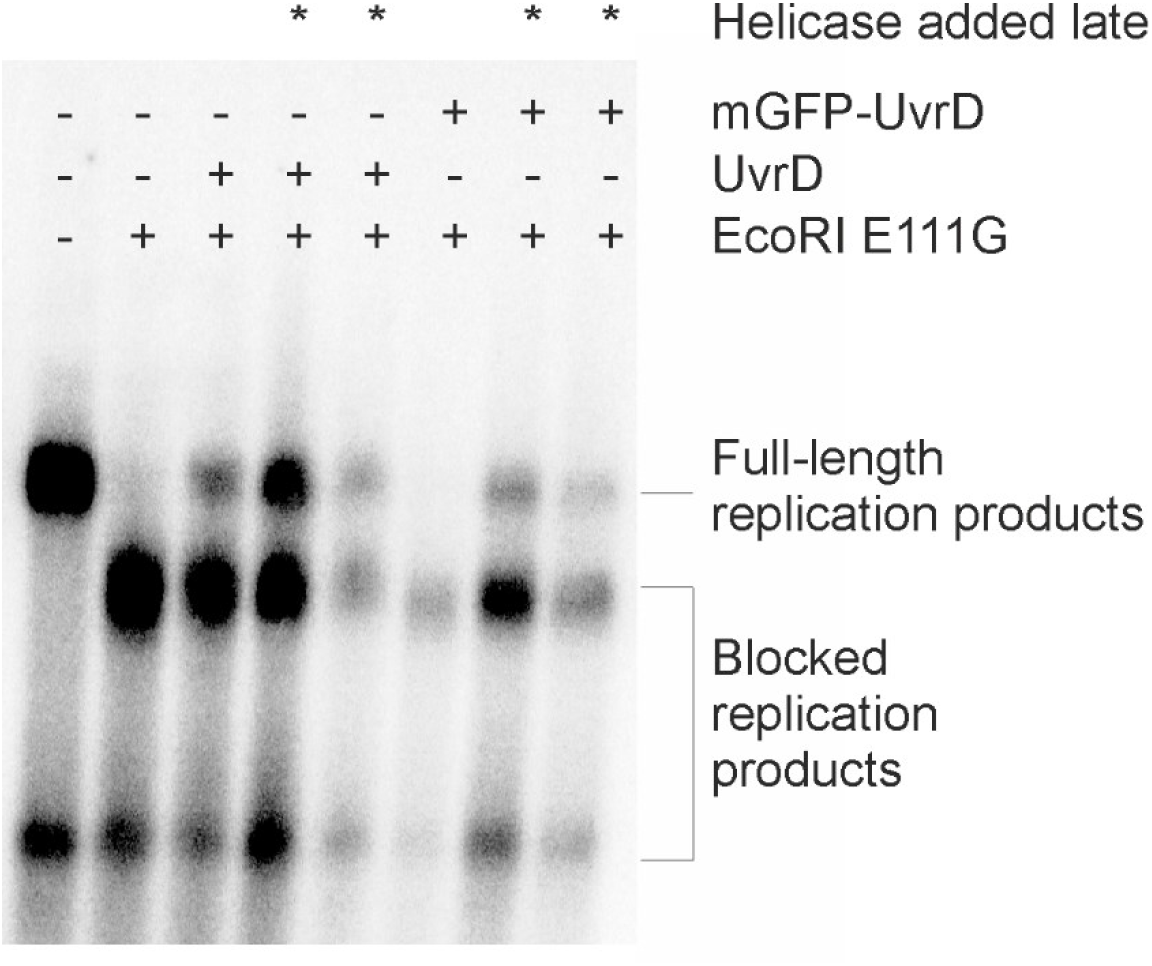
In vitro replication assay using mGFP-UvrD. Denaturing agarose gel of replication products formed from pPM872 (1) in the absence and presence of EcoRI E111G block, mGFP-UvrD and UvrD added pre-and post-collision (*) as indicated.

